# A near-complete genome assembly of the bearded dragon *Pogona vitticeps* provides insights into the origin of *Pogona* sex chromosomes

**DOI:** 10.1101/2024.09.05.611321

**Authors:** Qunfei Guo, Youliang Pan, Wei Dai, Fei Guo, Tao Zeng, Wanyi Chen, Yaping Mi, Yanshu Zhang, Shuaizhen Shi, Wei Jiang, Huimin Cai, Beiying Wu, Yang Zhou, Ying Wang, Chentao Yang, Xiao Shi, Xu Yan, Junyi Chen, Chongyang Cai, Jingnan Yang, Xun Xu, Ying Gu, Yuliang Dong, Qiye Li

## Abstract

**Background:** The agamid dragon lizard *Pogona vitticeps* is one of the most popular domesticated reptiles to be kept as pets worldwide. The capacity of breeding in captivity also makes it emerging as a model species for a range of scientific research, especially for the studies of sex chromosome origin and sex determination mechanisms.

**Results:** By leveraging the CycloneSEQ and DNBSEQ sequencing technologies, we conducted whole genome and long-range sequencing for a captive-bred ZZ male to construct a chromosome-scale reference genome for *P. vitticeps*. The new reference genome is ∼1.8 Gb in length, with a contig N50 of 202.5 Mb and all contigs anchored onto 16 chromosomes. Genome annotation assisted by long-read RNA sequencing greatly expanded the *P. vitticeps* lncRNA catalog. With the chromosome-scale genome, we were able to characterize the whole Z sex chromosome for the first time. We found that over 80% of the Z chromosome remains as pseudo-autosomal region (PAR) where recombination is not suppressed. The sexually differentiated region (SDR) is small and occupied mostly by transposons, yet it aggregates genes involved in male development, such as *AMH*, *AMHR2* and *BMPR1A*. Finally, by tracking the evolutionary origin and developmental expression of the SDR genes, we proposed a model for the origin of *P. vitticeps* sex chromosomes which considered the Z-linked *AMH* as the master sex-determining gene.

**Conclusions:** Our study provides novel insights into the sex chromosome origin and sex determination of this model lizard. The near-complete *P. vitticeps* reference genome will also benefit future study of amniote evolution and may facilitate genome-assisted breeding.

## Introduction

The central bearded dragon *Pogona vitticeps*, first described by German zoologist Ernst Ahl about a century ago, is an omnivorous lizard species belonging to the squamate family Agamidae [1]. In the wild, *P. vitticeps* has a natural distribution in the eastern and central Australia, where these lizards have adapted to the arid to semiarid habitats of the Australian inlands [2]. Yet, as a pet species, *P. vitticeps* is indeed commonly found around the world. It is one of the most popular reptile pets nowadays, owing to ease of breeding in captivity, presence of morphs varying in scale coloration, and docility to their owners [3]. And thanks to its easy accessibility and maintenance, *P. vitticeps* is also becoming a popular reptile model for a variety of research topics, such as amniote brain function and evolution [4, 5], chromosome rearrangement [6, 7], germline mutation rate [8], gene regulation of hibernation [9, 10], and especially, the mechanisms for sex chromosome origin and sex determination [11, 12], as it possesses an ZZ/ZW genetic sex determination system whereas the final sex is also influenced by environmental temperature during embryonic development [13].

While many studies involving *P. vitticeps* in the past decade tightly rely on its genome assembly and annotation, the current reference genome is still quite fragmented. This reference genome was constructed with a wild-caught ZZ male individual, based on Illumina short-read sequencing data generated from gradient libraries with insert size ranging from 250 bp to 40 kb [14]. Due to the limitation of short-read sequencing, the contig N50 of this assembly version is merely 35.5 kb in length, lag far behind the common standard of a reference genome (> 1 Mb) as proposed by the Earth BioGenome Project (EBP) [15] and the Vertebrate Genomes Project (VGP) [16] in recent years. Although this draft genome assembly was later scaffolded by different approaches that finally made ∼42 % of the genomic sequences anchored to chromosomes [6], the low anchoring percentage still limits its use in chromosome-scale investigation. For example, the sex chromosomes of *P. vitticeps* are well-known to be heteromorphic based on cytogenetic evidence, implying the existence of sequence divergence between Z and W due to recombination suppression [17]. However, the PAR and SDR of both sex chromosomes remain undefined so far, which in turn hamper the search for the master sex-determining gene.

Motivated by the increasing demand of *P. vitticeps* in academic research and pet industry, we decided to upgrade the genome assembly of this model lizard, via a combination of long- and short-read whole genome sequencing (WGS) as well as long-range sequencing technologies. The new *P. vitticeps* reference genome has a contig N50 of 202.5 Mb, with all contigs anchored onto 16 chromosomes and only few gaps remained. With this near complete chromosome-scale genome assembly, we fully characterized the Z sex chromosome and demarcated the PAR and SDR on Z, tracked the evolutionary origin and developmental expression of the Z-linked SDR genes, and proposed an alternative model for explaining the origin of sex chromosomes with the Z-linked *AMH* as the candidate of master sex-determining gene in *P. vitticeps*.

## Results

### The construction of a near-complete *P. vitticeps* genome

To reduce interference caused by allelic variation, all sequencing data for genome assembly were collected from a single captive-bred individual. The sex of this individual was verified as a ZZ male, as demonstrated by the anatomical presence of testes and the polymerase chain reaction (PCR) examination of sex-linked markers (Supplementary Fig. S1A). High molecular weight (HMW) DNA was extracted from the muscle and lung tissues, which was subjected to WGS with the CycloneSEQ [18] long-read and DNBSEQ short-read technologies, respectively (Supplementary Tables S1). In addition, we generated long-range sequencing data from the liver tissue by integrating chromatin conformation capture technique with CycloneSEQ (see Methods for details; hereafter referred as CycloneSEQ based Pore-C). *K*-mer analysis with the short-read data confirmed that *P. vitticeps* has a heterozygous diploid genome (heterozygosity ∼1.63%) with a haploid size of ∼1.7 Gb [14] (Supplementary Table S4; Supplementary Fig. S1B).

We obtained a total of ∼265 Gb CycloneSEQ WGS reads, with over a quarter (∼105 Gb) achieving a sequence length longer than 40 kb (Supplementary Table S1). As the 40 kb+ reads alone had already covered the *P. vitticeps* haploid genome for ∼62 times, fully satisfying the requirements for long read-based *de novo* assembly, we therefore assembled the 40 kb+ CycloneSEQ WGS reads using NextDenovo [19] in the first step (Fig. 1A; Supplementary Fig. S1C). This resulted in a 1.81 Gb primary assembly with merely 106 contigs. The contig N50 was 60.9 Mb, surpassing the majority of squamate genomes published to date (Supplementary Table S7). Before further scaffolding by long-range data, we removed false duplications in the primary assembly with Purge_Haplotigs [20], and corrected sequence errors with short-read data using NextPolish [21]. After polishing, a 1.80 Gb assembly with 74 contigs were scaffolded by ∼33X long-range CycloneSEQ based Pore-C data with YaHS [22]. The order and orientation of each contig was also manually examined in Juicebox [23] to avoid misplacements. The final chromatin contact map clearly sorted all the contigs into 16 chromosomes, including six macrochromosomes and 10 microchromosomes (Fig. 1B).

**Figure 1:**
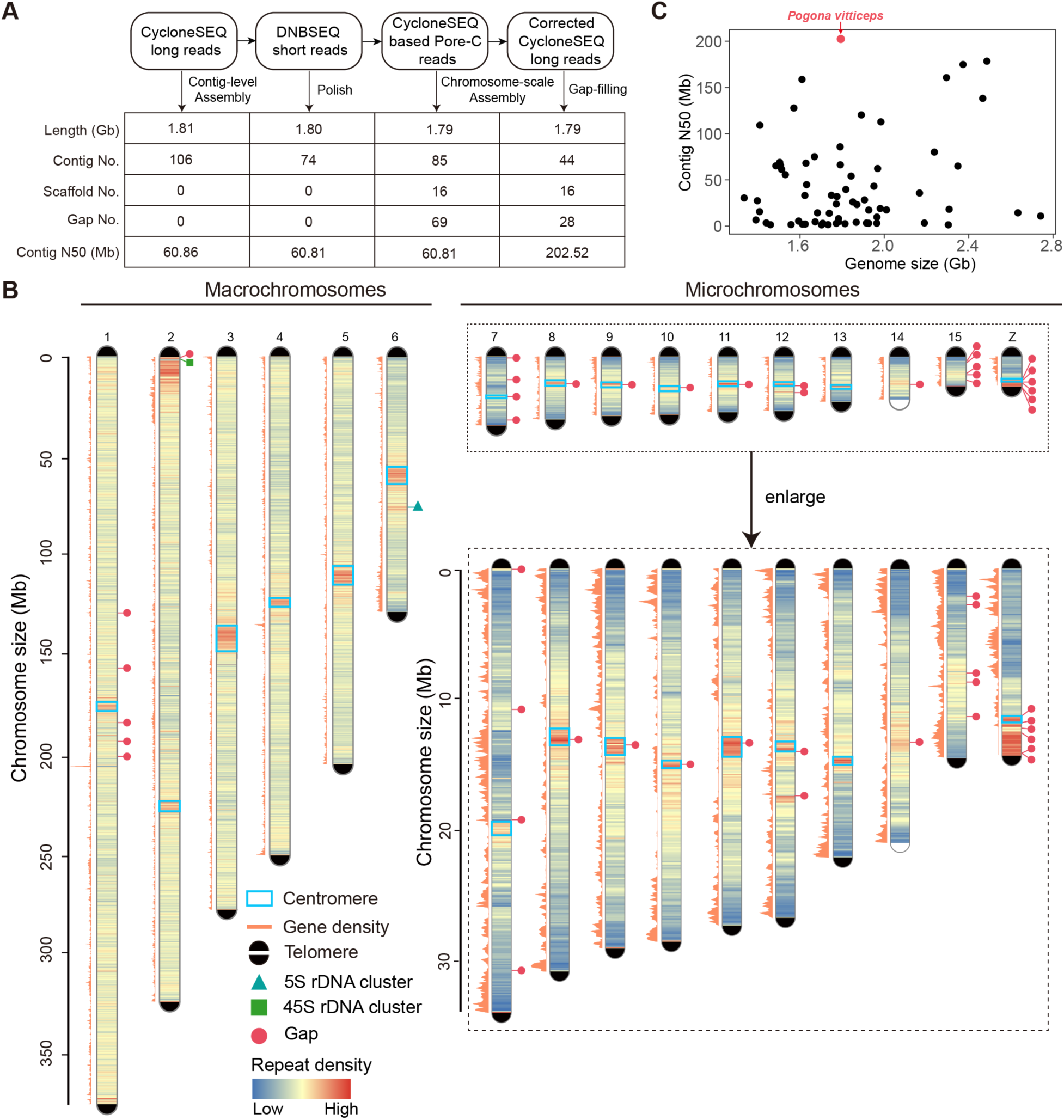
The near-complete genome assembly of *P. vitticeps*. (A) The CycloneSEQ long reads were initially utilized for *de novo* contig assembly. Thereafter, DNBSEQ short reads were employed to polish the assembled contigs. Subsequently, CycloneSEQ based Pore-C reads were implemented for scaffolding the contigs into chromosomal sequences. Finally, gaps were filled with corrected CycloneSEQ long reads. The table presents the genomic features for each step. (B) An overview of the genome features. The distribution of all repeat density and gene density were calculated in 100 kb windows. All the gaps in Pvit2024 were show in read circles. All chromosomes, except for chromosomes 14 and 15, had been identified with centromeric regions marked by light blue boxes. (C) The comparison of genome size and contig N50 between Pvit2024 and 66 publicly available squamate genomes assembled by long-read data.

To interrogate the integrity of chromosome ends, we next examined the presence of vertebrate telomeric repeat units (TRUs), i.e., (TTAGGG)_n_ [24]. Only eight of the 32 expected telomeres were presented in the initial chromosome-scale *P. vitticeps* assembly, probably due to the well-known challenge of assembling the highly repetitive telomeric regions [25] which was also evidenced by the prevalent absence of telomeric sequences in the chromosomal ends of most squamate genomes published so far (Supplementary Table S9). However, we were able to find many CycloneSEQ reads containing TRUs before assembling, suggesting that the missing of telomeres are mainly due to computational drawback rather than sequencing omission. We thus developed an in-house pipeline to conduct local telomere assembly with the TRU-containing reads and assign the assembled telomeres to their corresponding chromosome ends (see Methods for details). As a result, 31 of the 32 expected telomeres with a mean copy number of 1661 TRUs were patched to corresponding chromosome ends, except one on chromosome 14 (Fig. 1B).

After telomere repair, the chromosome-scale assembly still contains 69 unclosed gaps. These gaps were subjected to gap-filling with the corrected long reads (generated by NextDenovo during assembly) using TGS-GapCloser [26]. The correctness of each closed gap was further examined by long-read coverage with an in-house pipeline. This led to the solid closure of 41 gaps inside the chromosomes, and made four of the six macrochromosomes (chromosomes 3, 4, 5 and 6) as well as one microchromosome (chromosome 13) achieving telomere-to-telomere (T2T) gapless assembly. The remaining 28 gaps were mainly located in microchromosomes and co-localized with repeat-dense regions (Fig. 1B). Of note, a local region (0.5 to 11 Mb) with a particularly high density of repeats was clearly observed in almost all chromosomes, assumably corresponding to the position of centromeres (Fig. 1B; Supplementary Table S8).

In summary, the final *P. vitticeps* genome assembly (hereafter referred as Pvit2024) was 1.79 Gb in length, with all contigs anchored into 16 chromosomes, a final contig N50 of 202.5 Mb, and the presence of all but one of the 32 telomeres. Although there were still 28 inner gaps remaining unclosed, our Pvit2024 assembly outperformed all other squamate genomes reported so far in terms of continuity (Fig. 1C).

### Quality validation of the genome assembly

To assess the completeness of Pvit2024, we first aligned three sets of WGS reads to the Pvit2024 assembly, that is, the CycloneSEQ long reads and DNBSEQ short reads collected in this study as well as a set of Illumina short reads generated from another ZZ male individual that were not used for our genome assembly. All the three read sets revealed a mapping rate greater than 98% (Supplementary Table S5), and the aligned reads showed uniform coverage across the genome with only a few exceptions in repeat-dense regions (Fig. 2A). Secondly, we conducted Benchmarking Universal Single-Copy Orthologs (BUSCO) [27] and Compleasm [28] assessments for Pvit2024 with 7,480 Sauropsida conserved genes. The BUSCO and Compleasm complete score were 97.6% and 98.63%, respectively, comparable to or even higher than other squamate genomes assembled by long-read data (Supplementary Fig. S2A; Supplementary Table S7). The consensus quality value (QV) of Pvit2024 as estimated by short-read data was 36.4, corresponding to a single-base error rate of merely 0.0229%.

**Figure 2:**
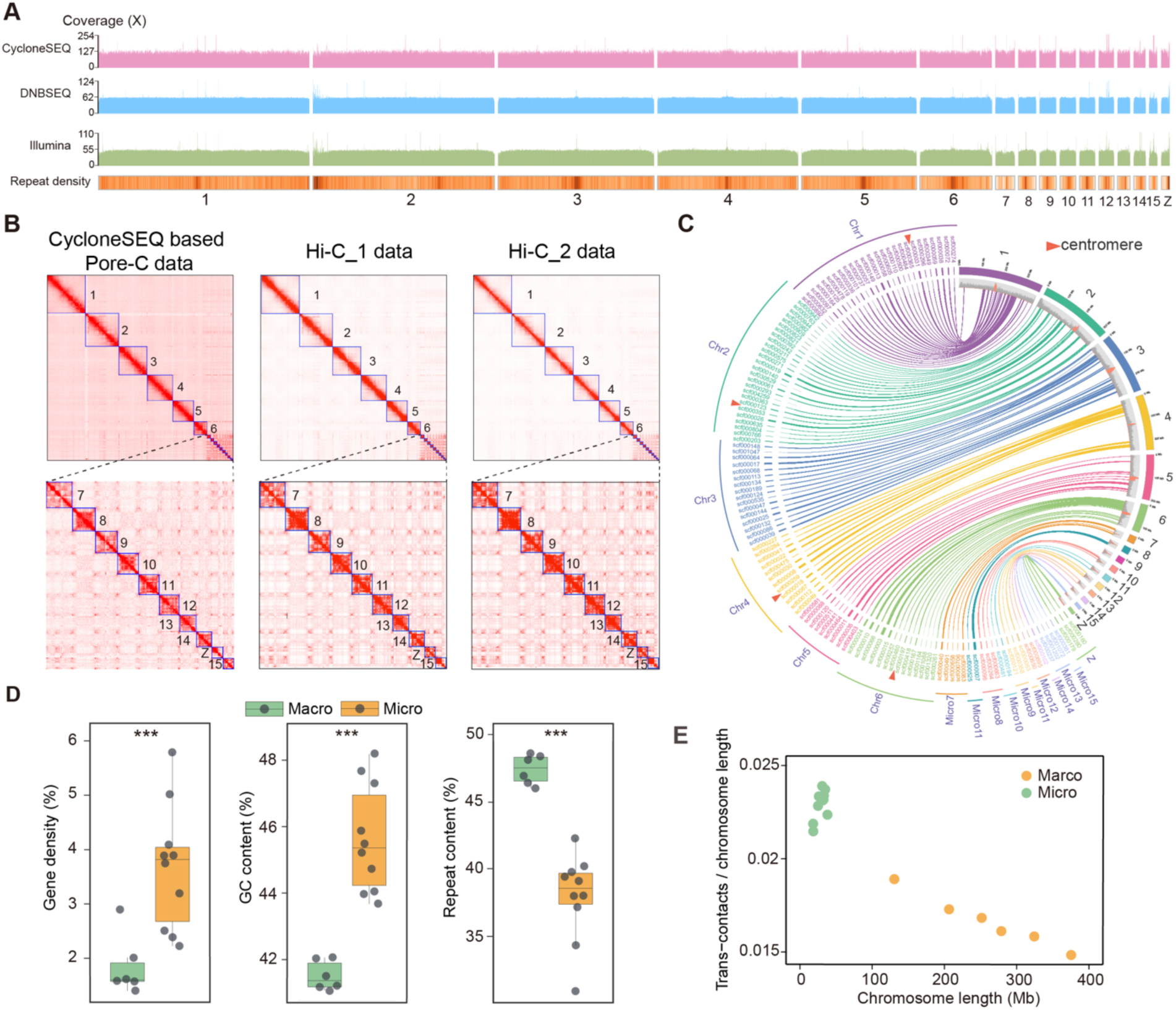
Quality validation of the genome assembly. (A) Genome-wide coverage based on various data was computed over 10 kb windows. Numbers on the y axis of each coverage track indicate the double average and average values of sequencing coverage for each track. Genomic data types are color coded. Pink: CycloneSEQ long-read WGS data. Light blue: DNBSEQ short-read WGS data. Light green: Illumina short-read WGS data. Additional tracks in the bottom panel show repeat density of Pvit2024. (B) The heatmap for one CycloneSEQ based Pore-C library and two traditional Hi-C libraries chromosomes contact matrices. (C) The circos plot illustrates genomic synteny, with Pvit2024 chromosomes on the right and scaffolds anchored to chromosomes by BACs from Deakin et al. The second right circle shows repeat density, red indicating centromeres. The red triangle marks the centromere position as determined by Deakin et al. (D) The comparison of gene density, GC content, and repeat content between macrochromosomes and microchromosomes. Asterisks indicate the significance of differences: *** represents p < 0.001. (E) Trans-contacts scaled by chromosome size for each chromosome were calculated by ∼33X CycloneSEQ based Pore-C data. Each dot represents one chromosome (yellow for macrochromosomes, green for microchromosomes).

Next, we assessed the accuracy of chromosomal sorting and scaffolding by different strategies. From a technical perspective, two traditional Hi-C libraries were constructed and sequenced with the DNBSEQ technology to serve as independent validations of the long-range information provided by CycloneSEQ based Pore-C. Both these two additional Hi-C contact maps consistently supported the scaffolding result of Pvit2024 (Fig. 2B). Besides, we aligned those chromosome-anchored sequences from Deakin et al. [6] to Pvit2024 and observed good agreements (i.e., sequences were mapped to expected position in expected order; Fig. 2C). Furthermore, Deakin et al. have also located the genomic regions harboring the centromeres in chromosomes 1, 2, 4 and 6 [6]. All these four centromere-containing regions showed perfect overlaps with the putative centromeric regions identified in Pvit2024 (Fig. 2C), supporting the feasibility of centromere positioning based on local repeat abundance in this species. From a biological perspective, the concurrent presence of macrochromosomes and microchromosomes is the hallmark of many vertebrate genomes [29]. According to the lengths of the assembled chromosomes, we could clearly define six of the sixteen Pvit2024 chromosomes as macrochromosomes and the remaining 10 as microchromosomes, in line with the reported karyotype of a male *P. vitticeps* (Supplementary Fig. S2C) [30]. In addition to chromosomal lengths, we also observed other recognized features that separate microchromosomes from macrochromosomes, including higher gene density, higher GC content, lower repeat content, and more frequent inter-chromosomal interaction (Fig. 2D, E; Supplementary Fig. S2B) [31, 32]. These multiple lines of evidence together highlight the reliability of the Pvit2024 chromosomal assembly.

### The new reference genome recovers ∼120 Mb missing sequences with numerous genes and regulatory elements

Compared with the previous reference genome (GCF_900067755.1/ pvi1.1) [14] (hereafter referred as Pvit2015), the contiguity of Pvit2024 was increased by over 5700 folds in terms of contig N50 (202.5 Mb vs 35.5 kb), and the number of scaffolds was remarkably reduced from 13,749 to 16 (Fig. 3A). Additionally, the genomic completeness as assessed by read alignments and BUSCO and Compleasm analyses all showed improvements (Fig. 3B, C).

**Figure 3:**
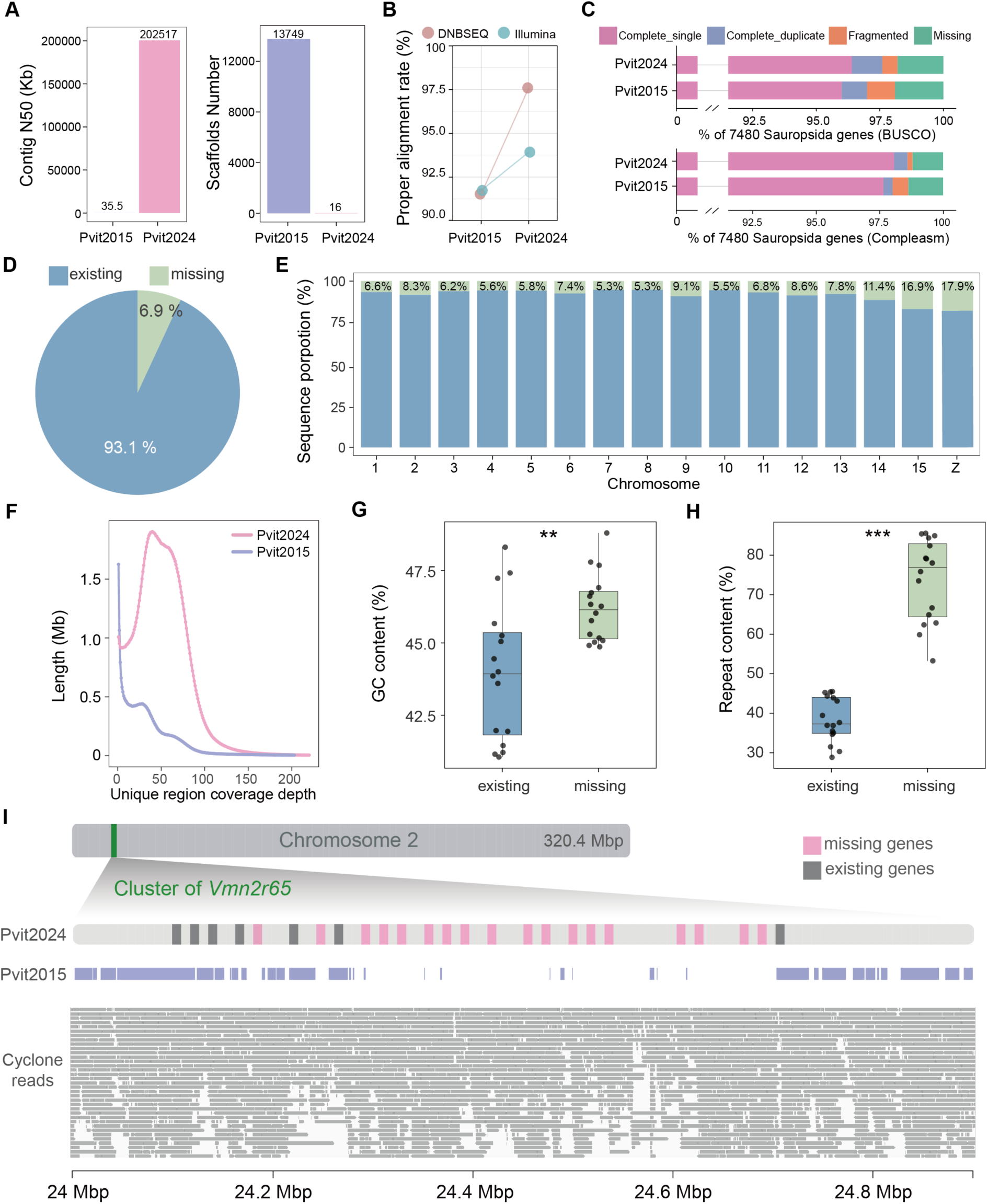
Comparison between Pvit2015 and Pvit2024. (A) The comparison of contig N50 and scaffold number. (B) The proper alignment rate of DNBSEQ and Illumina WGS data. (C) Gene completeness assessment of Pvit2015 and Pvit2024 based on BUSCO and Compleasm analyses. (D) Proportion of sequences existing and missing in the Pvit2015 assembly compared to the Pvit2024. (E) The missing rate of Pvit2015 assembly across chromosomes in Pvit2024 is depicted, with blue and green bars representing the existing and missing ratios of Pvit2015, respectively. (F) Base coverage depth distribution of unique sequences in Pvit2015 and Pvit2024, the DNBSEQ genomics data were used. (G and H) Comparison of GC content (G) and repeat element content (H) between existing and missing sequences in Pvit2015 across each chromosome. T-test analysis of variance; \**P*<0.05, \*\**P*<0.01, \*\*\**P*<0.001. (I) Distribution of *Vmn2r65* gene family along the chromosome, determined using ChromoMap of R packages. The blue block show sequence of Pvit2015 that can be aligned back to Pvit2024. The grey strip of bottom shows the coverage of CycloneSEQ reads.

To further uncover the differences between the two genome assemblies, we conducted reciprocal whole-genome alignment to identify missing sequences in one assembly relative to the other. Briefly, we defined genomic sequences in one assembly with no alignment to the other one by Winnowmap2 [33] and Minimap2 [34] as missing sequences in the former one. In this way, 124 Mb of the Pvit2024 sequences were detected as missing in Pvit2015, accounting for 6.9% of the Pvit2024 assembly size and affecting all the 16 chromosomes in a different degree (Fig. 3D, E; Supplementary Fig. S3A). Read coverages of these Pvit2015-missing regions were uniform and close to that of genome average, indicating that they are not assembly errors (Fig. 3F; Supplementary Fig. S3H). Conversely, only 36 Mb of Pvit2015 sequences were not aligned to Pvit2024, and the read coverage of these regions were quite low, suggesting that these Pvit2015-speficic sequences likely resulted from assembly errors (Fig. 3F).

We next focused on the annotation of the 124 Mb sequences that were missing in prior assembly, especially for the possibility of carrying functional genes or regulatory elements. The missing sequences revealed higher GC content than genome average, and most missing sequences (∼73.6%) was annotated as repetitive elements in Pvit2024, reminding the advantage of the long-read sequencing to sequence through high GC content regions and repeat regions (Fig. 3G, H; Supplementary Fig. S3B) [35]. Nevertheless, it is noteworthy that these Pvit2015-missing sequences also harbored the intact copies of up to 576 protein-coding and 312 lncRNA genes, namely, they are completely absent in prior assembly (Supplementary Fig. S3C, D). A representative example was the *Vmn2r65* (Vomeronasal 2 Receptor) gene family, of which 25 copies were located on a tandem array in chromosome 2 of Pvit2024 (Fig. 3I); only seven copies of *Vmn2r65* were identified and distributed on 6 distinct scaffolds in Pvit2015. In addition, the missing sequences in Pvit2015 also led to partial absence of one or more exons for up to 1,319 protein-coding and 1,651 lncRNA genes (Supplementary Fig. S3C, D), which may affect the accuracy of expression quantification by RNA sequencing. In terms of regulatory elements, we first examined the completeness of gene promoters, the region upstream of genes where the RNA polymerase binds to initiate transcription [36] and found that 2,504 (11.5%) protein-coding and 1,281 (8.1%) lncRNA genes contain >500 bp missing sequences in their putative promoter regions, respectively. We then examined the CpG islands, which are widespread in vertebrate genomes and play important roles in transcriptional regulation [37]. Consistent with the difficulty of short-read technology to sequence through high GC regions, we observed that 32,864 of the annotated 216,126 (15.2%) CpG islands were completely missing and 7,361 (3.4%) were partial missing in prior Pvit2015 assembly (Supplementary Fig. S3E, F, G).

Taken together, our genome-wide comparative analysis indicated a notable proportion (6.9%) of genomic sequences missing in the prior *P. vitticeps* reference genome. These missing sequences tend to be GC-rich and repeat-rich, and more importantly, they contain numerous genes and regulatory regions that are not accessible based on prior reference genome.

### Improving genome annotation by long-read RNA sequencing

Besides genome assembly, comprehensive and accurate annotation of gene model is the necessity to realize the value of a reference genome [38]. To facilitate a comprehensive gene annotation for Pvit2024, we performed high-depth long-read RNA sequencing (RNA-seq) for eight different tissues (brain, eye, heart, kidney, liver, lung, muscle, and testis) with the CycloneSEQ technology (Supplementary Table S2). In the meanwhile, we also generated paired-end short-read RNA-seq data for these eight tissues with the DNBSEQ technology (Supplementary Table S3). All the long-read and short-read RNA-seq data were generated from the same individual as that used for genome sequencing in this study.

With the abundant RNA-seq data, we first conducted protein-coding gene annotation with a combination of transcriptomic, homologous and *ab initio* prediction evidence, and obtained a total of 21,783 non-redundant protein-coding gene models (Supplementary Table S11). Up to 92% (20,002) of the protein-coding loci are supported by RNA-seq signal (TPM > 5) in at least one tissue. BUSCO assessment with the Sauropsida conserved genes revealed a complete score of 97.5%, consistent with that estimated for Pvit2024 genome assembly and slightly higher than the commonly used NCBI annotation (96.6%) and Ensembl annotation (91.7%) based on Pvit2015 (Fig. 4A).

**Figure 4:**
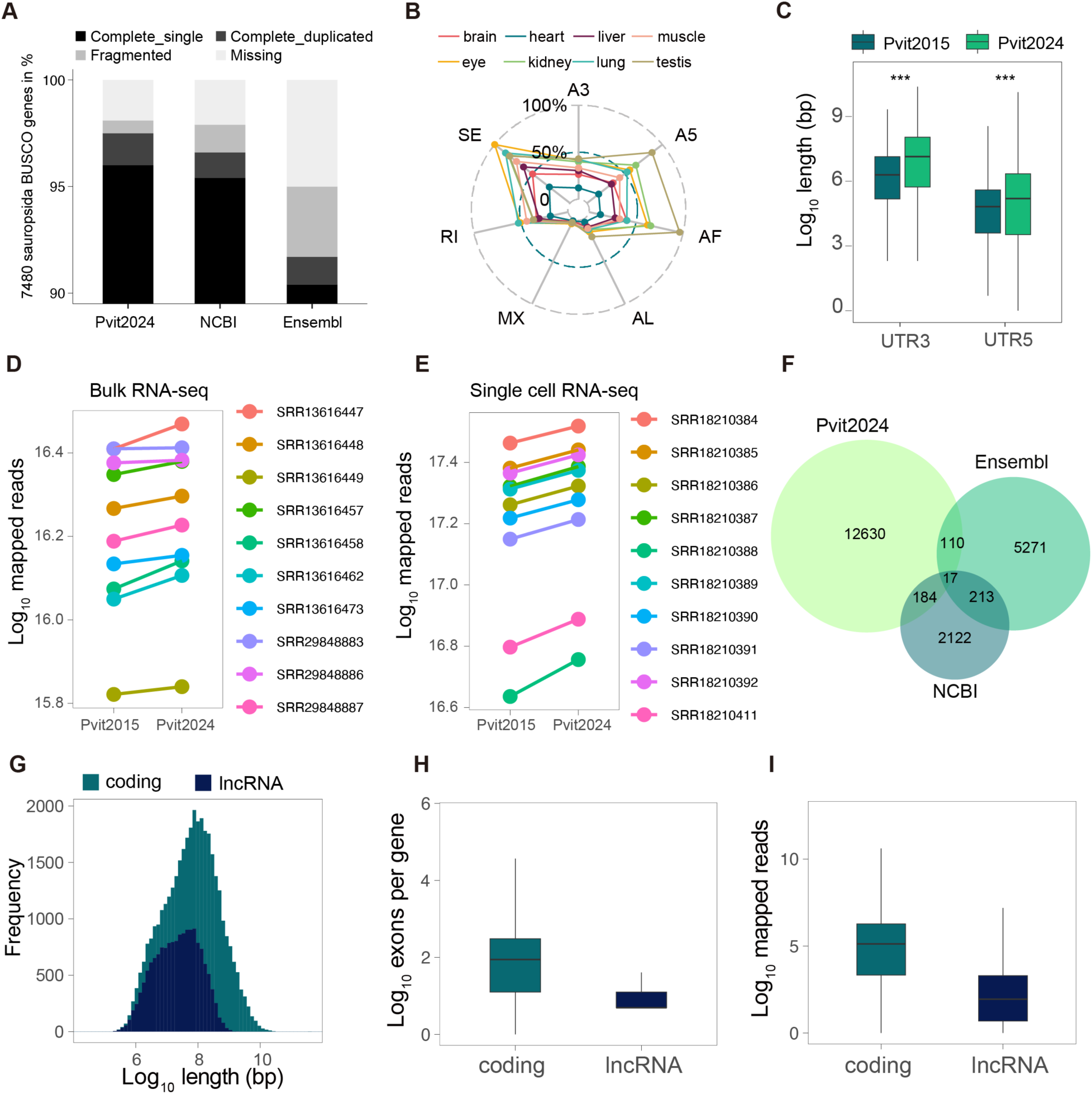
Enhancement of the genome annotation for *P. vitticeps*. (A) Completeness of Sauropsida BUSCO genes in annotations produced by Pvit2024, NCBI (Pvit2015), and Ensembl (Pvit2015). (B) Radar chart showing relative splicing abundance of eight tissues for each event type (SE–skipping exon; RI–retained intron; MX-mutually exclusive exons; A5–alternative 5’ splice site; A3–alternative 3’ splice site; AF-alternative first exons; AL-alternative last exons). Normalization here employs the maximum abundance value of splicing events as the denominator. (C) UTRs length distributions between the Pvit2024 and NCBI, Pvit2024 has longer than NCBI. T-test analysis of variance; \**P*<0.05, \*\**P*<0.01, \*\*\**P*<0.001. (D and E) Connected dot-line showing the variance of mapping reads in Pvit2024 and NCBI RefSeq annotations for bulk (D) and single-cell (E) transcriptomes. (F) Venn diagram showing lncRNA counts for Pvit2024, Ensembl and NCBI RefSeq annotations. (G) Histogram shows the distribution of gene length (exon regions) for coding gene and lncRNA. (H) Distribution of the exon counts per gene in coding gene and lncRNA. (I) Distribution of the read counts for transcriptomes in coding genes and lncRNAs.

Then, we leveraged the long-read RNA-seq data to refine isoform and UTR annotations for the protein-coding genes. After strict filtering steps to exclude artificial and low-quality sequences, we finally assigned 53,272 transcripts to 21,783 protein-coding loci, with a mean of 2.5 isoforms detected per gene. Up to 97% of the splicing junctions derived from the assigned transcripts were also supported by short-read alignments, further supporting the reliability of these transcript models. In addition, all the major types of alternative splicing events could be identified in all the eight tissues, with exon skipping and alternative first exon usage being the most dominant events (Fig. 4B). By discriminating coding regions from non-coding parts, we could identify the UTRs for most transcripts, with 17,110 (31.1%) having a 5’-UTR longer than 30 bp and 19,161 (34.8%) having a 3’-UTR longer 50 bp. Of note, while the lengths of coding regions were comparable between our and the NCBI/Ensembl annotations, the 5’-UTRs and 3’-UTRs in our annotation were significantly longer (Fig. 4C), corroborating a more accurate definition of transcription boundaries owing to the assistance of long-read RNA-seq. To evaluate the effect of the new annotation on future RNA-seq studies, we calculated the read mapping rate and number of expressed genes with a bulk and a single-cell RNA-seq dataset that were not used for gene annotation in this study. Both datasets revealed a significant improvement on both metrics, especially for the single-cell dataset (Fig. 4D, E).

Long non-coding RNAs (lncRNAs) are another important class of RNA molecules that engage in numerous biological processes [39], yet the lncRNA repertoire is understudied in squamate reptiles. By identifying transcripts that are longer than 200 nt and lack coding potential, we annotated 13,269 high-confidence lncRNAs in Pvit2024 (Fig. 4F). When compared with the protein-coding genes, the *P. vitticeps* lncRNAs were generally shorter in length, have fewer exons, and were expressed at lower levels (Fig. 4G, H, I), consistent with previous observations in other organisms [40]. Interestingly, we found that many lncRNAs were expressed specifically in one or a few tissues, especially the testis (Supplementary Fig. S4). The large amount of tissue-specific lncRNAs in uncovered in *P. vitticeps* implied their potential importance in squamate biology that worth further attention.

### The demarcation of PAR and SDR along the *P. vitticeps* Z sex chromosome

The identity of *P. vitticeps* Z chromosome was ascertained by mapping known Z-linked scaffolds identified by Deakin et al. [6] to our Pvit2024 assembly. Specifically, all the Z-linked scaffolds were aligned to a ∼14.3 Mb microchromosome in expected order, spanning a continuous region of ∼8.2 Mb on the first half of this microchromosome (Fig. 5A; Supplementary Fig. S5A). The uncovered portion comprised ∼6.1 Mb genomic sequences that were newly identified to be Z-linked, accounting for ∼43% of the whole Z chromosome (Fig. 5A). These newly identified Z-linked sequences harbored 47 protein-coding genes and 65 lncRNA genes that are either missing or unknown to be Z-linked in prior Pvit2015 assembly (Supplementary Table S12). The overall Z chromosome showed an uneven distribution of genomic elements, with an increase of repeat content accompanied by depletion of protein-coding genes toward the chromosomal end that enriched newly anchored sequences (Fig. 5B).

**Figure 5:**
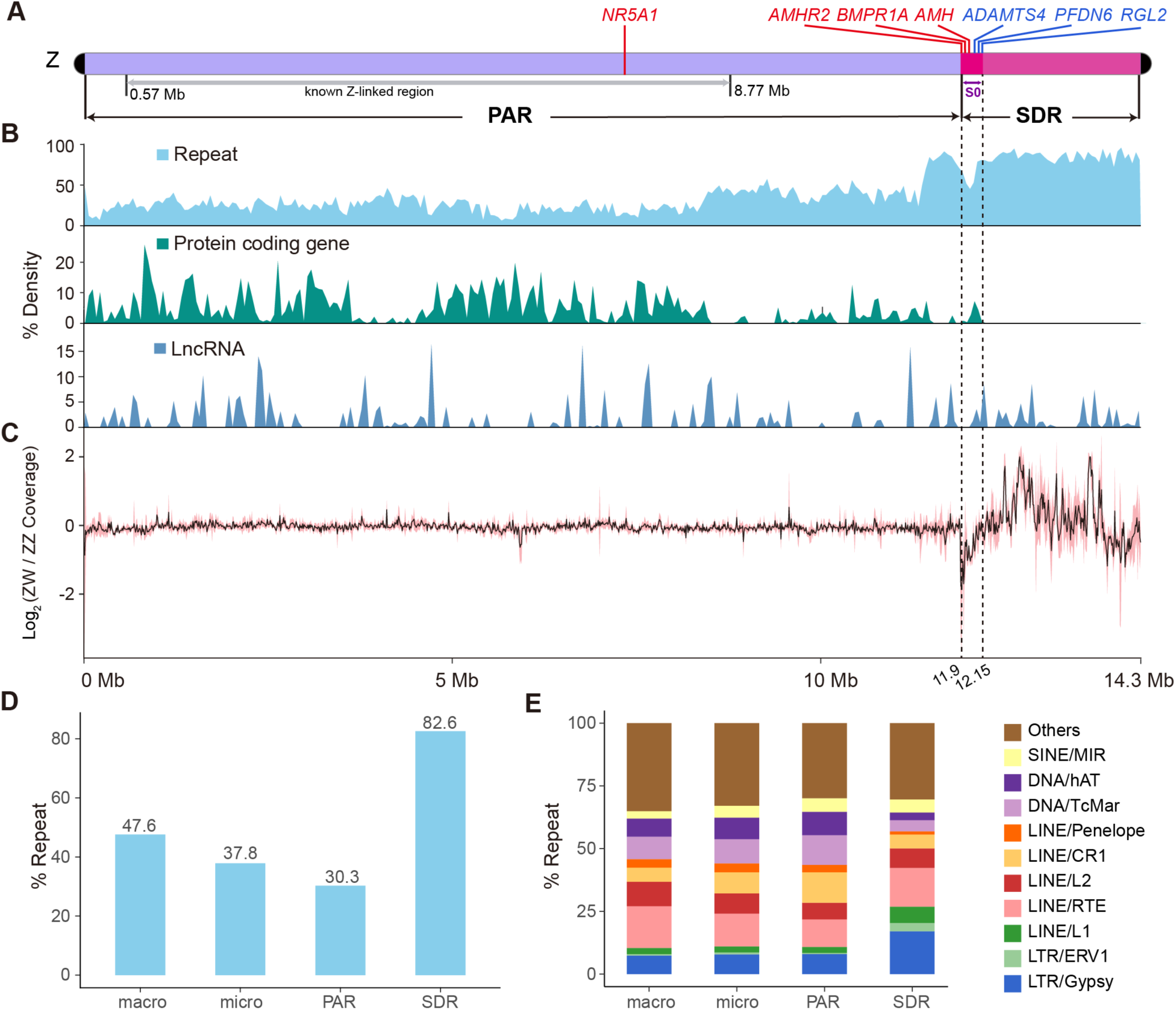
Z sex chromosome identification and the composition of PAR and SDR. (A) The Z chromosome can be divided into two regions: PAR and SDR. Sex-determination genes are marked in red, while pseudogenes are in bule. (B) Repeat density, protein coding gene density, and lncRNA density are calculated in 50 kb windows. (C) The distribution shows the ratio of ZW to ZZ read coverage in 10 kb windows. The depth (black line) is calculated as the average ratio of read coverage for ZW females to ZZ males between each pair of ZW females and ZZ males, based on the WGS data from two ZW females and four ZZ males. The pink areas are the confidence intervals for each window. (D) Repeat content for macrochromosomes, microchromosomes, PAR region, and SDR region. (E) Relative proportions for ten most abundant transposons families and other elements.

We next attempted to demarcate the SDR, where Z-W sequence divergence is accumulated, and the PAR, where Z and W remains identical, by comparing WGS read coverage between ZW and ZZ individuals. In principle, a PAR is expected to display comparable read coverage in both genders, while a SDR would exhibit halved coverage in ZW females relative to ZZ males [41]. Guided by this basic idea, we analyzed the WGS data collected from two ZW females and four ZZ males. Up to 83% (∼11.9 Mb) of the Z chromosome displayed a comparable read coverage in both genders, suggesting that the vast majority of Z remains as PAR that persists recombination. Nevertheless, there was a ∼250 kb region exhibiting roughly halved coverage in ZW females relative to ZZ males (Fig. 5C), fulfilling the expectation of an evolutionarily old stratum (hereafter referred as S0) where Z and W have been substantially diverged. For the region ranging from the downstream of S0 to the chromosome end, the read coverage in ZW females was even higher than that in ZZ males (Fig. 5C). However, this region lacked protein-coding genes and instead was occupied exclusively by repetitive elements (Fig. 5B). We speculated that this unusual coverage pattern was attributed to the much higher abundance of repeats accumulated in the W counterpart of this region, as previous cytogenetic studies has suggested that the short arm of W is much longer than Z [42]. If so, this downstream region of S0 (∼2.18 Mb in length) might also represent a stratum that was fully degenerated due to transposon invasion, and therefore, we tentatively grouped it into SDR in this study.

Together, we could conservatively define a large, continuous PAR and a small SDR on the *P. vitticeps* Z chromosome, with the PAR-SDR boundary delimited at ∼12 Mb in coordinate (Fig. 5A). Compared with PAR, SDR was apparently characterized by a remarkably high repeat content (>80%) (Fig. 5D). However, the repetitive element composition of SDR was generally similar to those of PAR and other chromosomes, although several classes of transposons such as LTR/Gypsy, LTR/ERV1 and LINE/L1 were relatively more abundant in SDR (Fig. 5E).

### The evolutionary origin and developmental expression of the Z-linked SDR genes

Although being predominantly occupied by repetitive elements, we found that the *P. vitticeps* SDR carried three intact protein-coding genes (*AMHR2*, *BMPR1A* and *AMH*) and three pseudogenes (*ADAMTS4*, *PFDN6* and *RGL2*), of which all located in a ∼150 kb region within S0 (Fig. 5A). Notably, the three intact genes are all well-known players in driving sexual differentiation, namely, *AMH* (anti-Mullerian hormone) [43] and its two receptors, *AMHR2* (anti-Mullerian hormone receptor type 2) [44] and *BMPR1A* (bone morphogenetic protein receptor type 1A) [45]. It is also noteworthy that each of these six SDR genes had a counterpart (i.e., paralog) on autosomes, while all other investigated lizards encoded only one copy of each in their genomes (Fig. 6A), indicating that lineage-specific gene duplication events occurred in *P. vitticeps* (and probably in other closely related agamid lizards as well). This was also supported by gene phylogenetic analyses, which preferentially clustered the *P. vitticeps* paralogs together among the proteins from multiple species (Fig. 6B, Supplementary Fig. S6A). In contrast, while the PAR maintained up to 334 protein-coding genes, only 21 of them (6.3%) had paralogs in autosomes. Hereafter, we designated the six SDR genes as *AMHR2-Z*, *BMPR1A-Z*, *AMH-Z*, *ADAMTS4-Z*, *PFDN6-Z* and *RGL2-Z*, and their autosomal counterparts as *AMHR2-A*, *BMPR1A-A*, *AMH-A*, *ADAMTS4-A*, *PFDN6-A* and *RGL2-A*, respectively (Fig. 6C).

**Figure 6:**
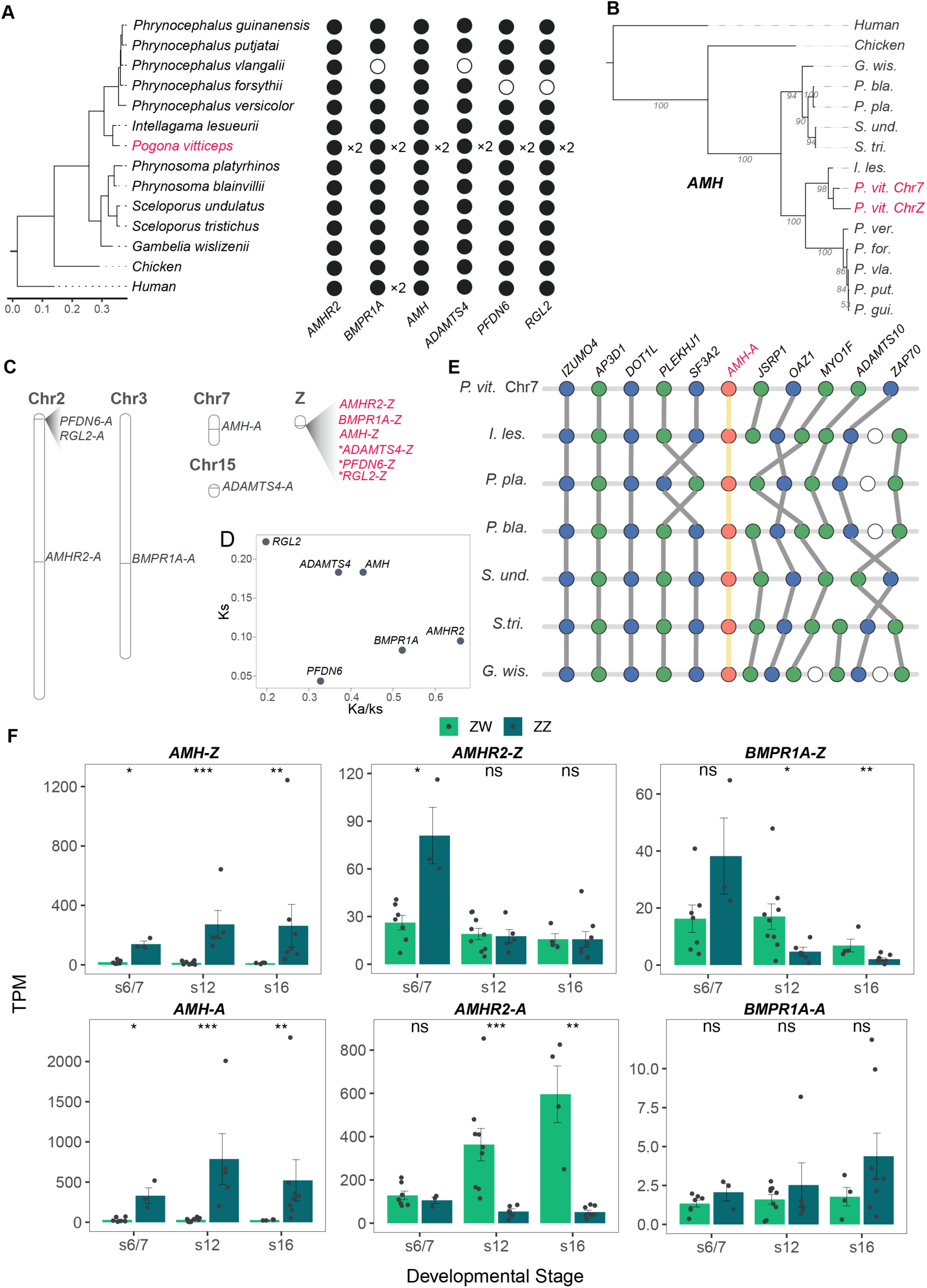
Z-linked SDR genes. (A) Copy number of SDR genes in cross-species. A dot (black) indicated a copy, digital followed by an “×” indicated copy number, and circles indicated absence. (B) Phylogenetic relationships of *AMH* genes across different species. The pink font showing the two copies of *AMH* in *P. vitticeps*. (C) Ideogram showing the gene model of SDR genes and their autosomal counterparts in the Pvit2024 assembly. Pseudogenes were indicated with an asterisk. “-A” indicated autosomal genes, while “-Z” indicated sex-linked genes. (D) Ka/Ks analysis of SDR genes showing order of gene occurrence in Z-linked SDR. (E) Synteny relationships of *AMH-A* and its flanking 5 genes in closely related species. *AMH* is highlighted in orange, and surrounding genes are shown in grey. (F) Differential expression of SDR genes between ZZ and ZW during developmental stages. Benjamini-Hochberg analysis of variance: ns (not significant), \**P*<0.05, \*\**P*<0.01, \*\*\**P*<0.001.

We next asked, for these six paralog pairs in *P. vitticeps*, whether the SDR or the autosomal copy was the original copy that gave rise to the other one. By examining gene synteny information across species, we found that none of the SDR copies could be deemed to be original, namely, they all arose from the duplication of their autosomal counterparts (Fig. 6D, Supplementary Fig. S6B). But it is noted that *PFDN6* and *RGL2* were probably duplicated as a whole, because their gene order as well as the transcriptional direction in SDR maintained the same as their original copies in chromosome 2 (Fig. 6C; Supplementary Fig. S6B). We then estimated the synonymous nucleotide substitution rate (Ks) between the paralog pairs to date the order of their translocation into SDR. As expected, the pseudogenes displayed the largest Ks, probably due to relaxed selection. But it is interesting that, among the three intact genes in SDR, *AMH* displayed the largest Ks, suggesting that *AMH* is likely among the earliest ones to be integrated into the SDR (Fig. 6D).

The master sex-determining gene that initiates sex differentiation remains unknown in *P. vitticeps*, yet it is expected to be a gene that locates in SDR and reveals differential expression between ZZ male and ZW female at an early stage of gonad differentiation. Therefore, we examined the expression dynamics of the SDR genes along gonadal development with the RNA-seq data from Wagner et al. [46]. This dataset comprised gonadal transcriptomes of both genders collected at three embryonic developmental stages (stage 6/7, stage 12 and 16). In *P. vitticeps*, stage 6/7 represents the earliest stage at which a consolidated gonad is recognizable and begins to differentiate, stage 12 represents an early stage of differentiation, while the gonads at stage 16 have been fully differentiated [13, 46]. Of note, all the three intact SDR genes (*AMH-Z*, *AMHR2-Z* and *BMPR1A-Z*) were identified as differentially expressed genes (DEGs) between genders at one or more developmental stages (DESeq2 FDR < 0.05; Fig. 6F), while none of the pseudogenes and lncRNAs in SDR were significant DEGs (Supplementary Fig. S6C). In addition, we found that *AMH-Z* (FDR = 0.00005; FC = 5.54) and its potential receptor *AMHR2-Z* (FDR = 0.05; FC = 2.22) manifested a significant male-biased expression pattern at stage 6/7, the earliest developmental stage examined. Moreover, *AMH-Z* maintained and even amplified the differential expression pattern throughout gonadal differentiation, reinforcing its central role in driving male development in vertebrates. Together with its early integration into SDR, *AHM-Z* could be considered as a strong candidate of the master sex-determining gene in *P. vitticeps*.

Meanwhile, we also examined the expression of the autosomal paralogs of the SDR genes, to explore whether the expression pattern has been diverged after gene duplication. Interestingly, *AMH-A* displayed almost the same expression pattern as *AMH-Z*, maintaining a high expression level in males while repressing its expression in females throughout embryonic gonadal development (Fig. 6F). However, the expression dynamics of *AMHR2-A* and *BMPR1A-A* were apparently diverged from that of their SDR counterparts. Specifically, while *AMHR2-Z* displayed transient male-biased expression at stage 6/7, *AMHR2-A* was not differentially expressed at stage 6/7 but turned to be significantly female-biased at stages 12 and 16 due to its upregulation in ZW females. In terms of *BMPR1A*, the expression of *BMPR1A-Z* was a bit male-biased at stage 6/7, but the following sharp decrease of *BMPR1A-Z* expression in ZZ males made it turn to be significantly female-biased at later stages; however, *BMPR1A-A* did not show significant sex-biased expression throughout development, although a slight trend of male-biased expression was observed (Fig. 6F).

## Discussion

### The first chromosome-scale reference genome for Amphibolurinae (Squamata: Agamidae)

Squamate reptiles are a species-rich clade of amniote vertebrates that have adapted to most terrestrial ecosystems [47]. With over 12,000 extant species that make up a significant part of the vertebrate tree of life, reference genomes for squamates are particularly scarce when compared with other amniote clades such as mammals and birds [48]. Specifically, according to the latest data deposited in NCBI (accessed in July 2024), only 104 squamate species have a genome assembly with contig N50 over 30 kb, in sharp contrast with ∼630 for birds and ∼730 for mammals. This unevenness has drawn the concern of the biodiversity genomics community who call for more attention for this neglected vertebrate group in the genomic era [15, 16]. Thanks to technological advances in long-read sequencing, constructing a near-complete reference genome has become feasible for many organisms, especially for those able to supply sufficient HMW DNA from a single individual, such as the reptiles as showcased in this study. By leveraging a newly released long-read sequencing technique called CycloneSEQ, we upgraded the genome assembly of the model lizard *P. vitticeps* to one of the most continuous squamate genomes so far. This new *P. vitticeps* genome represents the first chromosome-scale reference genome for the Amphibolurinae subfamily of the agamid lizards (Agamidae), highlighting its potential value in evolutionary genomics studies. With this near-complete *P. vitticeps* reference genome, we also provided novel insights into the sex chromosome origin and sex determination of this species.

### AMH signaling as an upstream driver of sexual differentiation in *P. vitticeps*

*P. vitticeps* possesses a ZZ male/ZW female genetic sex determination (GSD) system. However, the default sexual fate of ZZ embryos can be overwritten by high incubation temperature, leading to sex reversal of ZZ males to ZZ females [49]. The capacity of a ZZ embryo to develop as a normal female without the help of W chromosome suggests that sexual fate is most likely determined by a dosage-sensitive gene on the Z chromosome [12]. Therefore, the achievement of a complete Z chromosome sequence represents a particularly important beginning for disclosing the sex-determining cascade in *P. vitticeps*. Although there are several unclosed gaps remained in repeat-rich regions, our Pvit2024 Z chromosomal assembly is approaching T2T level as indicated by the presence of telomeric sequences on both chromosomal ends. Additionally, the Pvit2024 Z has almost doubled the known Z-linked sequences, from 8.3 Mb to 14.3 Mb. More importantly, the newly anchored Z-linked sequences carry the whole SDR, a region that hopefully harbors the master sex-determining gene.

The Z/W-resided *nr5a1*, which encodes the steroidogenic factor 1 (SF1) required for male development, was once regarded as the most promising sex-determining candidate in *P. vitticeps* [50]. However, the location of *nr5a1* in PAR as uncovered in our study has almost ruled out it as the master sex-determining gene. Instead, the aggregation of *AMH* signaling related genes (i.e., *AHM*, *AHMR2* and *BMPR1A*) in the SDR is of particular interest. *AMH*, which encodes a hormone of the transforming growth factor-β (TGF-β) superfamily, is critical for testis development and has been identified as a master sex-determining gene in a growing number of vertebrates, including several lineages of teleost fish [51] and monotreme mammals [52]. In *P. vitticeps*, we found that the SDR-resided *AMH* (*AHM-Z*) was highly expressed in all examined ZZ embryos and maintained over 5-fold expression difference between ZZ and ZW embryos since the beginning of gonadal differentiation (stage 6/7). This prominent male-biased expression pattern suggests *AMH-Z* as a strong sex-determining candidate, which deserves further functional experimental validation in future studies.

AMH signals through binding to its receptors, such as the type II receptor encoded by *AMHR2* and the type I receptor encoded by *BMPR1A* [45]. Therefore, our finding of transient up-regulation of the SDR-resided *AMHR2* and *BMPR1A* in ZZ embryos at stage 6/7 is particularly notable. We hypothesize that such a transient up-regulation of the *AHM* receptors may serve as an amplifier of the *AHM* signaling cascade at the bipotential gonads, which is essential for tipping the sex differentiation network towards male development at this critical time window. Another notable finding is the up-regulation of the autosome-resided *AMH* (*AMH-A*) since stage 6/7. Given the high protein similarity of *AHM-Z* and *AMH-A* that implies their functional conservation, this parallel up-regulation of *AMH-A* may serve as another booster of AMH signaling during early sex differentiation. Collectively, all this evidence points to one conclusion, that is, the AMH signaling is essential and likely serve as an upstream signal driving *P. vitticeps* sex differentiation. Such pattern of duplicated gene functioning in upstream sex determination have previously been identified mainly in fish [53]. And interestingly, all these duplications, including our current finding in the *Pogona* lizard, seems to direct gonad development into testis.

However, it is also notable that the gonadal transcriptome profiles of ZZ and ZW embryos has been diverged at stage 6/7 [46]. This suggests that the initial elevation of AMH signaling likely occurs even earlier, before stage 6/7. Interestingly, in teleost fish Patagonian pejerrey (*Odontesthes hatcheri*) a convergent gene duplication event was found as *AMH* was duplicated onto the Y chromosome and may also serve as the candidate sex determinant gene, indicative by an earlier expression than the autosomal *AMH* [54], Future studies are recommended to focus on when and where (e.g., what cell types) the AMH signaling is initially diverged between ZZ and ZW embryos.

### A proposed model for the origin and evolution of *P. vitticeps* sex chromosomes

The origin of the *P. vitticeps* ZW microchromosomes is presumably associated with the fusion of a fragment derived from chromosome 2, as both the Z and W chromosomes share homology with the terminal region of the long arm of chromosome 2 (chr2qter) as revealed by physical mapping of a BAC clone (Pv151P16) [55]. Our result also supports this homology according to the finding of an additional copy of the chr2qter-resided *PFDN6-RGL2* syntenic block in SDR (Fig. 6C). However, we also found that the homology with chr2qter is limited to the terminal region of the *P. vitticeps* Z chromosome which lacks intact protein-coding genes, thus arguing the role of the chr2qter fusion event in driving ZW formation. Indeed, the fusion of chr2qter to *P. vitticeps* ZW microchromosomes seems to occur early during Amphibolurinae evolution, probably before the differentiation of Z and W, because the concurrent mapping of Pv151P16 to chr2qter and a pair of microchromosomes are also observed in other Amphibolurinae species, including GSD species with homomorphic sex chromosomes and TSD species without sex chromosomes [55].

However, it is notable that the *Ks* of *AMH*-Z is higher than that of *PFDN6-Z* and almost comparable to that of *RGL2-Z*, despite that *PFDN6-Z* and *RGL2-Z* have become pseudogenes (Fig. 6C). This suggests that the integration of *AMH* and the fusion of chr2qter to the proto-sex chromosomes might occur within a very narrow time window. We thus propose that the duplication and translocation of the autosomal *AMH* to the proto-sex chromosomes might represent the real milestone in triggering the formation of Z and W chromosomes in the ancestor of *P. vitticeps* and its sister taxa. After *AMH* integration, the subsequent aggregation of other male-biased genes including *AMHR2* and *BMPR1A* further consolidated the role of the proto-Z in sex determination and promoted ZW differentiation. We anticipate that chromosome-scale genome assemblies from other closely related dragon lizards will provide crucial evidence for testing this hypothesis.

## Methods

### Sample collection

This study was performed in accordance with the guideline of the national and organizational stipulation. An adult lizard *Pogona vitticeps* in captivity was collected at Zhejiang University, under the permit ZJU20240342. This lizard was euthanized after anesthetized by diethyl ether. All the tissue samples were frozen via liquid nitrogen and stored at −80℃ refrigerator immediately after dissection. Muscle and lung tissues were selected for genome sequencing, which included CycloneSEQ long-read WGS sequencing, DNBSEQ short-read WGS sequencing, CycloneSEQ based Pore-C sequencing, and Hi-C sequencing. Furthermore, eight tissue samples, including liver, lung, eye, muscle, testis, kidney, brain, and heart, were subjected for transcriptome sequencing.

### Genome and Transcriptome sequencing

The High-molecular-weight genomic DNA was extracted using the cetyl trimethylammonium bromide method (see “Genomic DNA CTAB extraction” for details) from muscle and lung samples. Eight tissue samples were used for RNA extraction with TRlzol reagent (Invitrogen, USA) following the manufacturer’s guidelines. The isolated RNA was then fragmented into 200-400 bp, and then reverse-transcribed to cDNA for library preparation. Quantity and quality of the genome’s DNA and RNA were assessed by pulsed field gel electrophoresis, Qubit 3.0 (Invitrogen, USA) and Qseq 400 (Bioptic, China). A total of 12 short-insert paired-end (PE) libraries (4 for genomic DNA and 8 for cDNA) were constructed and sequenced on DNBSEQ platform (MGI, Shenzhen), with the manufacturer’s instructions. The long-read libraries were constructed as CycloneSEQ library and sequenced via CycloneSEQ WuTong02 platform (see “CycloneSEQ library preparation and sequencing” for details). For the Hi-C library construction, the muscle sample was crosslinked with formaldehyde and 2 Hi-C libraries were constructed by using the dpnII restriction endonuclease. The libraries were sequenced on the DNBSEQ platform with 150-bp paired-end sequencing strategy.

### Genomic DNA CTAB extraction

Preheat the CTAB lysate in a water bath. Collect 20-30 mg of fresh or frozen animal tissue, freeze in liquid nitrogen. Grind tissue using mortar and pestle in the presence of liquid nitrogen until finely ground. Transfer the ground animal tissue to 2 mL polypropylene centrifuge tubes. Add 1 mL of CTAB lysis buffer and 50 µL protease K (20 mg/mL). swirl and mix, incubate at 50 ℃ (with shaking) for 1 h. Cool the tube to 37 °C. Add 20 µL RNase A (10 mg/mL) for every 1 mL of lysis buffer, mix by inversion and incubate for 10 min. Add equal volume phenol-chloroform-isoamyl alcohol (25:24:1 ratio) with pH > 7.8, mix by inversion and spin at 5000 rpm for 10 minutes in a tabletop centrifuge at room temperature (RT). Transfer top aqueous solution to new 2 mL centrifuge tubes using wide-bore pipette tip. Add equal volume chloroform-isoamyl alcohol (24:1 ratio), mix by inversion and spin at 5000 rpm for 10 minutes in a tabletop centrifuge at RT. Transfer top aqueous solution to new 1.5 mL centrifuge tubes using wide-bore pipette tip. Add 2/3 volume of isoamyl alcohol and mix by inversion to form an emulsion, centrifuge at 5000 rpm for 2 minutes at 4°C. Discard the supernatant. Add 1mL 75% ethanol and mix slowly to resuspend the precipitation. Centrifuge at 5,000 ×g for 2 min at 4°C and discard the supernatant. Repeat the previous step. Discard the remaining supernatant, dry at RT for 3 min. Dissolve in 200-400 µL TE Buffer and incubate at 37 °C for 1 h, incubate at RT overnight. Store the extracted DNA at −80 °C.

### CycloneSEQ library preparation and sequencing

The preparation and sequencing of CycloneSEQ library was proceeded following the manufacturer’s guidelines. Each sample, comprising 2 μg of input DNA (≥23 ng/μL), was initially diluted with nuclease-free water to a total volume of 44 μL, followed by mixing with 6 μL DNA repair buffer 1, 3 μL DNA repair buffer 2, 3 μL DNA repair enzyme 1, and 4 μL DNA repair enzyme 2. The mixes were then incubated in a thermocycler through the following steps: 10 min at 20℃, 10 min at 65℃, and held at 4℃. After incubation, the mixes were purified using a 1.0x volume of DNA clean beads, and DNAs were eluted with 60 μL of nuclease-free water. Next, the purified end-repaired samples were mixed with 10 μL sequencing adaptors, 25 μL 4x ligation buffer, 10 μL DNA ligase, and 2.5 μL nuclease-free water, before incubating at 25℃ for 30 minutes to complete the adaptor ligation. The ligated products were again purified with a volume of 1.0x DNA clean beads and long fragment wash buffer was applied to gently resuspend the beads. After removing the supernatant, the libraries were recovered into 17 μL of elution buffer and quantified on a Qubit fluorometer. Each prepared library was sequenced on the CycloneSEQ WuTong02 platform according to the protocol.

### CycloneSEQ based Pore-C sequencing

About 100 mg tissues were homogenized with pestle and washed with 2000 μL 1X PBS (Phosphate-Buffered Saline). After being spun down, the tissues were fixed by 10ml of 1% formaldehyde 1X PBS solution and incubated for 10 minutes at room temperature (RT). To quench formaldehyde crosslinking, 527 μL of 2.5 M glycine was added and incubated for 5 minutes at RT, followed by 10 minutes on ice. Fixed tissues were pelleted by centrifugation at 2000 ×g for 10 minutes at 4 °C. The pellet was resuspended in 550 μL of ice-cold lysis buffer (10 mM Tris-HCl pH 8.0, 10 mM NaCl, 0.2% Igepal-CA630, 50 μL of protease inhibitor cocktail) and incubated on ice for 15 minutes. Nuclei were pelleted at 2000 ×g for 10 minutes at 4 °C and the supernatant was discarded. The pellet was resuspended in 200 μL of the prepared chilled 1.5X digestion reaction buffer and was pelleted by centrifugation at 2000 ×g for 10 minutes at 4 °C. The pellet was resuspended in 300 μL of 1.5X digestion reaction buffer and added 33.5 μl 1% SDS, followed by incubate in a thermomixer at 300 rpm at 65°C for 10 minutes. The reaction was stopped by adding 37.5 μl of 10% (v/v) Triton X-100. Then, 45 μL of 10 U/μL NlaIII restriction enzyme (NEB, R0125L) and 34μL water were added, and the sample was rotated at 37 °C for 18 hours. NlaIII was then heat-inactivated at 65 °C for 20 minutes. A total of 550 μL of ligation master mix was added: 100 μL of 10× NEB T4 DNA ligase buffer with 10 mM ATP (NEB, B0202), 10 μL of 10 mg/mL BSA (Thermo Fisher, AM2616), 50 μL of 400 U/μL T4 DNA Ligase (NEB, M0202), and 390 μL of water. The reactions were rotated at RT for 5 hours. Then, the sample was added a master mix (100 μl of 10% SDS, 500 μL of 20% tween-20, 100 μl of 20 mg/ml proteinase K, 300 μL of water) and incubated at 56 °C for 18 hours. The sample was extracted with equal volume of phenol: chloroform: isoamyl alcohol (25:24:1) and precipitated with 0.7 X isopropanol. DNA was size selected (>1.5 kb) using DNA Clean beads and constructed to CycloneSEQ library.

### Sex identification

Organ anatomical examination and sex-linked markers analysis were employed for sex identification in *P. vitticeps*. For the anatomical assessment, testes were dissected from the individual. Molecular analysis of sex-linked markers was conducted using a protocol adapted from Holleley et al. [11], Genotypic sexing was performed utilizing two PCR primers: H2, GCCCATATCTCACTAGTTCCCCTCC; F, CAGTTCCTTCTACCTGGGAGTGC, which was flanking two W-chromosome-specific deletions, measuring 150 base pairs and 14 base pairs, respectively. PCR was conducted using Platinum High-Fidelity ReadyMix(2x) (GCATbio), with a range of primer concentrations and genomic DNA quantities to establish experimental groups, and a no-genomic DNA control group. Cycling conditions were 95℃ for 5 min; (95℃ for 20 s, 70∼65℃ for 20 s, 72℃ for 1 min) ×10 cycles with annealing temperature decreased 0.5℃ per cycle; (95℃ for 20 s, 65℃ for 20 s, 72℃ for 1 min) ×30 cycles; 72℃ for 10 min. The PCR products were resolved on a 1.5% agarose gel and visualized using GelStain Blue (Transgen). The presence of two bands indicated ZW individuals, while a single band confirmed ZZ individuals.

### Genome survey

The DNBSEQ short-read WGS data was cleaned by SOAPnuke v1.5.6 (RRID: SCR_015025) [56] to exclude reads characterized by low quality and the presence of adapter sequences and poly-N regions with parameters -Q 2 -G -d −l 20 -q 0.2 −5 1 -t 5,0,5,0. To accurately assess genomic characteristics, including genome size and heterozygosity rate, all clean data were adopted *k*-mer based methods by using Jellyfish and GenomeScope tools. The haploid genome size was estimated according to *k*-mer analysis frequency distributions generated by Jellyfish v2.2.6 (RRID: SCR_005491) [57] using a series of *k* value (19, 21, 23, 25, 27) with the -C setting, which was calculated as the number of effective *k*-mers (ie. total *k*-mer – erroneous *k*-mer) divided by the homozygous peak depth. The rate of heterozygosity was estimated by GenomeScope v2.0.0 (RRID: SCR_017014) [58] with the *k*-mer frequency distributions generated by Jellyfish as inputs.

### Genome assembly

The initial process was to retain CycloneSEQ long reads longer than 40 kb, the chimeric reads were removed using Yacrd v1.0.0 [59] with parameters -c 5 -n 0.6. Cleaned reads were then assembled into contigs using NextDenovo v2.5.0 (RRID:SCR_025033) [19] with the following parameters: read_cutoff = 35k, genome_size = 1.7g. In response to the high heterozygosity detected in the initial assembly, DNBSEQ short reads were adopted to remove heterozygous contigs by purge_haplotigs v1.1.2 (RRID:SCR_017616) [20] with parameters −l 10 -m 53 -h 120. Next, two rounds of polishing were performed using NextPolish v1.4.1 (RRID:SCR_025232) [21] with recommended parameters, utilizing DNBSEQ short reads. Subsequently, CycloneSEQ based Pore-C reads were used as the main body to anchor the assembly to pseudo-chromosomes, while two additional libraries of Hi-C reads served as controls to assess the reliability of the anchoring. The specific process of anchoring was as follows: CycloneSEQ based Pore-C reads were first aligned to the contig-level genome using wf-pore-c v1.1.0 (https://github.com/epi2me-labs/wf-pore-c) with default parameters. The unaligned fragments were subsequently filtered out from the wf-pore-c output file ’null.ns.bam’ using the command ’samtools view -F 4 -bh’. Adjacent fragment pairs were then extracted from the filtered BAM file into BED format with a custom script (https://github.com/guoqunfei/Pvit_T2T). In parallel, two additional libraries of Hi-C reads were aligned using Chromap v0.2.3-r407 [60] with default parameters. The alignment results were converted from the .sam format to a .bed file using bedtools v2.29.2 [61]. Both sets of BED files, derived from CycloneSEQ based Pore-C and Hi-C reads respectively, were utilized to assemble contigs into scaffolds using YaHS v1.2a.1 [22] with the parameters –no-contig-ec –no-scaffold-ec. Following the scaffolding process, we proceeded to generate a .hic file for further manual examination and curation using JuiceBox (JBAT) [23], adhering to the guidelines provided at https://github.com/c-zhou/yahs.

To avoid telomeric repeat sequences being incorrectly trimmed by the assembler, by using known vertebrate six base telomere repeats (‘TTAGGG’) as a sequence query, we developed an in-house pipeline to correct telomeric regions as follows: First, we extracted those reads from CycloneSEQ WGS raw reads containing at least 100 consecutive copies of the telomere units. Then, all of these reads were aligned to the assembly using minimap2 v2.23-r1116-dirty (RRID:SCR_018550) [34] with the parameter: -x map-ont. Subsequently, based on the alignment results, for each chromosome’s two ends, the read with the highest alignment quality and the largest number of copies of the telomere unit was selected and used to replace the corresponding end. (The specific implementation code can be seen at https://github.com/guoqunfei/Pvit_T2T).

After telomere repair, unclosed gaps in the chromosome-scale assembly were initially filled using TGS-GapCloser v1.2.1 (RRID:SCR_017633) [26] with the corrected long reads (generated by NextDenovo during assembly). The correctness of each closed gap was further examined by long-read coverage with an in-house pipeline as follows: the 5 kb of upstream and downstream regions were extracted and aligned to the corrected long read to identify any alignment. These alignments were further visualized and manually examined using IGV. Only alignments with proper size and consistent orientation without conflicting alignments were used for the purpose of gap filling within the genomic assembly.

Finally, the assembled genome was evaluated using BUSCO v5.7.1 (RRID:SCR_015008) [27], Compleasm v0.2.6 [28], Merqury v1.3 (RRID:SCR_015811) [62] using default parameters. Short reads were aligned to the genome using BWA-MEM v0.7.17-r1198-dirty (RRID:SCR_012940) [63], and SAMtools v1.15.1 (RRID:SCR_002164) [64] were counted as properly paired. Long reads were mapped to the genome using minimap2 v2.23-r1116-dirty (RRID:SCR_015008) [34] with the parameter: -ax map-ont.

### Reconstruction of NOR 45S rDNA cluster

According to the comprehensive cytogenetic map, *P. vitticeps* has an active nucleolus organizer region (NOR) rich in 45S rDNA cluster, comprising of 18S, 5.8S, and 28S rRNAs, at the sub-telomeric region of Chromosome 2 [42]. However, the sequencing depth of the NOR region, identified as the 45S rDNA-rich region near the telomere on Chr2 from the *de novo* assembly of whole genome, was significantly higher than that of adjacent regions. Additionally, the length of the NOR region was much shorter than those found in closely related species. These findings indicated that the NOR region was not fully assembled. To address this, we reconstructed the NOR 45S rDNA cluster. Firstly, the 45S rDNA sequence from NOR region was used to identify all CycloneSEQ reads containing the 45S rDNA sequence feature. Secondly, these reads were assembled using NextDenovo v2.5.0 [19] with the following parameters: read_cutoff = 1k, genome_size = 0.005g, minimap2_options_raw = -I 6G --step 2 --dual=yes -t 4 -x ava-ont -k 17 - w 17 --minlen 2000 –maxhan1 5000. Finally, we obtained a 282.2 kb NOR region rich in 45S rDNA cluster and replaced this segment with the genomic NOR position where the assembly had collapsed.

### Repetitive element annotation

We annotated the lizard whole-genome repeat sequences based on *de novo* predictions and homology annotations. For *de novo*, the RepeatModeler v2.0.4 (RRID:SCR_015027) [65] was used to identification the custom repeats from the assembly sequence, with subsequent annotation and masking using RepeatMasker v4.1.5 (RRID:SCR_015027) [66]. For homology annotations, the homologous repeat elements in the genome were identified and classified using RepeatMasker. The transposable element proteins were searched based on RepeatProteinMask database (http://www.repeatmasker.org) that part of RepeatMasker, and the tandem repeats were extracted using TRF v4.09 [67] via ab initio prediction.

### Protein coding gene annotation

We integrated three types of evidence for predicting protein-coding genes: transcriptome data, homology evidence, and ab initio prediction. For the transcriptome evidence, we first cleaned the DNBSEQ RNA-seq data using SOAPnuke v1.5.6 (RRID:SCR_015978) [56] with the command: filter -n 0.03 −l 20 -q 0.3 -p 1 -Q 2 -G −5 1 -t 10,0,10,0 -E 70. Next, we aligned the cleaned short reads to the genome using HISAT2 v2.2.1 (RRID:SCR_015530) [68] with the –dta option. Finally, the transcripts were reconstructed using StringTie2 v2.2.3 (RRID:SCR_016323) [69] and coding sequences were predicted with TransDecoder v5.7.1 (RRID:SCR:015534) (https://github.com/TransDecoder/TransDecoder). For the ab initio prediction, we randomly selected 700 high-quality genes from the transcriptome predictions for training and used AUGUSTUS v3.4.0 (RRID:SCR_008417) [70] for the ab initio annotation of coding genes. For homology-based prediction, homologous data from five closely related species including *Ahaetulla prasina* (GCF_028640845.1), *Furcifer pardalis* (GCA_030440675.1), *Hemicordylus capensis* (GCF_027244095.1), *Rhineura floridana* (GCF_030035675.1), *and Zootoca vivipara* (GCF_963506605.1), were collected by NCBI RefSeq. To sum up, we integrated the above three types of evidence and short reads splice information data as input to GeMoMa v1.9 (RRID:SCR_017646) [71] workflow for a comprehensive prediction of coding genes. For the annotation results obtained from the GeMoMa, we further filtered based on the repeat sequence annotation file, removing predictions suspected to be repeat elements, ultimately obtaining the final protein coding gene annotation set.

### Isoform detection by long reads

The process of CycloneSEQ long-read RNA-seq data was as follows: we filtered the TSO (template switch oligo sequence)/RTP (reverse transcription primer) and split chimeric reads based on artificial sequence. The clean data was mapping to assembly using winnowmap2 v2.03 [33] with the command: *winnowmap -u b -K 4G -G 135k -ax splice ref fastq*. We then reconstructed transcripts based on the reference annotation using Isoquant v3.5.0 [72] with the command: *python3 isoquant.py –gene_quantification all –reference genome.fa –data_type nanopore –genedb genes.gtf –threads 16 –count_exons –bam input.bam –label sample*. Subsequently, we merged the transcripts from eight tissues, and eliminate redundancy based on an all-exon overlap threshold of greater than 0.95 between isoforms. We then performed TransDecoder v5.7.1 to predict the coding potential of all transcripts using the *–single_best_only* option. Finally, isoforms of coding gene were assigned to the correspond loci.

### LncRNA annotation

LncRNA identification also based on long sequencing reads of CycloneSEQ RNA-seq. The remaining transcripts assembled by Isoquant from the isoform detection steps, which filtering by length (≥200 nt), were used as the foundational dataset for lncRNA detection. We then utilized two tools of FEELnc v0.2.1 [73] and CPC2 v1.0.1 (Coding Potential Calculator, RRID:SCR_002764) [74] to filter potential coding genes. The candidates of lncRNA were obtained by taking the intersection of the results from both tools. We further mapped the corresponding DNBSEQ sequencing data from the same library with CycloneSEQ RNA-seq using HISAT2 v2.2.1 for alignment to the assembly. Expression detection of candidate lncRNAs was performed using FeatureCounts v2.0.1 (RRID:SCR_012919) [75]. We ultimately considered lncRNAs with TPM > 1 to be true and reliable.

### *De novo* prediction for tRNA and rRNA

Two types of noncoding RNAs were predicted, namely transfer RNAs (tRNAs) and ribosomal RNAs (rRNAs). To identify putative tRNA genes in the *P. vitticeps* genome, we employed tRNAscan-SE v2.0.10 (RRID:SCR_008637) [76] with parameters optimized for eukaryotic genomes. Subsequently, we filtered out “pseudo” and “undet” from the output generated tRNAscan-SE, and extract high-confidence tRNA genes from the BED file, producing a refined tRNA set. For rRNA prediction, we employed the barrnap v0.9 (RRID:SCR_015995) (https://github.com/tseemann/barrnap) program to identify rRNA genes in the genome, using the following command: barrnap –quiet –kingdom euk genome.fa –threads 20.

### Chromosome contact analyses

In the above scaffolding procedure, the .hic files were derived from the processing of CycloneSEQ based Pore-C reads and Hi-C reads. Subsequently, the interaction contacts within the .hic files were binned to construct the genome-wide interaction matrix at resolutions of 5 kb, 10 kb, 100 kb-, 500 kb, 100 kb, 500 kb and 1Mb. Following this, the ICE (iterative correction and eigenvector decomposition) normalization was then employed to normalize the interaction matrix. Then, cis-contacts and trans-contacts for each 100-kb window were calculated using the 100-kb normalized interaction matrix.

### Missing sequence identification

We compare the Pvit2024 genome assembly with a previous (Pvit2015) assembly generated using Illumina sequencing. The Pvit2015 assembly was downloaded from NCBI RefSeq by searching for GCF_900067755.1. We excluded the mitochondrial genome from the assembly to prevent misalignment between mitochondrial and nuclear genomes. We then aligned the Pvit2015 assembly to Pvit2024 ref by minimap2 v2.23-r1116-dirty (RRID:SCR_018550) [34] with the following command: *minimap2 -ax asm20 -k14 -K 8G –secondary=no -s 80 -t 16 Pivt2015 Pvit2024*, and with paftools to obtain the aligned regions by minimap2. Additionally, we also performed alignments using the Winnowmap2 tool, which requires the creation of a database with Meryl v1.4.1 [62]. The database was created using the following commands: *meryl count k=19 output merylDB Pvit2024* and *meryl print greater-than distinct=0.9998 merylDB > repetitive_k19.txt*. The alignment was then executed with the command: *winnowmap -t 16 -W repetitive_k19.txt -K 8G –secondary=no -ax asm20 -s 80 Pvit2024 Pvit2015*. Similarly, we used paftools to obtain the alignment files for the two genomes. We further utilized a Python scripts (https://github.com/guoqunfei/Pvit_T2T) to extract unaligned regions from the alignment files. The final set of missing regions was determined by taking the intersection of the results obtained from both methods.

### Missing sequences in genomic elements

We identified the proportions of coding genes, lncRNAs, and promoter within the missing regions based on coordinate overlap information. To identify missing coding genes and lncRNA, we considered a locus span and exon region with an overlap greater than 0.8 with the missing region as a missing gene in the Pvit2015 assembly. In contrast, an overlap between 0.1 and 0.8 was classified as an incomplete gene. Additionally, for the identification of promoters, we defined the 2000 bp upstream region of coding genes and lncRNAs as potential regulatory regions. An overlap greater than 0.7 between these regions and the missing region was considered a missing promoter, while an overlap between 0.1 and 0.7 was classified as an incomplete promoter.

### Calculation of GC content and repeat content

GC content was determined by calculating the total number of Gs and Cs divided by the length of the given coordinates, excluding ambiguous nucleotides (N), using a custom Python script. Repeat content was assessed based on pre-annotated repeat files, with a Python script used to extract the coordinates of repeat regions and count the number of repeat nucleotides.

### Identification of S0 region

To identify regions with differential sequencing coverage between the Z and W chromosomes of *P. vitticeps*, two ZW female WGS data and four ZZ male WGS data were aligned to the assembled genome using BWA-MEM v0.7.17-r1198-dirty (RRID:SCR_012940) [63] (Supplementary Table S13). Depth of coverage was extracted using SAMtools v1.15.1 (RRID:SCR_002164) [64]. Median depth was calculated using a non-overlapping sliding window of 10 kb.

### Searches for genes in SDR regions of closely related species

The genome assemblies of all closely related species, including *Gambelia wislizenii* (GCA_030847615.1), *Intellagama lesueurii* (GCA_037013535.1), *Phrynocephalus forsythia* (GCA_029282475.1), *Phrynocephalus guinanensis* (GCA_037367245.1), *Phrynocephalus putjatai* (GCA_037367255.1), *Phrynocephalus versicolor* (GCA_023846285.1), *Phrynocephalus vlangalii* (GCA_037367305.1), *Phrynosoma blainvillii* (GCA_026167975.1), *Phrynosoma platyrhinos* (GCA_020142125.1), *Sceloporus tristichus* (GCA_016801415.1), *Sceloporus undulatus* (GCA_019175285.1), as well as the outgroup species *Gallus gallus* (GCF_016699485.2) and *Homo sapiens* (GCF_009914755.1), were downloaded from NCBI RefSeq. We used the protein sequences of *AMHR2*, *BMPR1A*, *AMH*, *ADAMTS4*, *PFDN6*, and *RGL2* from human, chicken, and green anole, which were downloaded from Ensembl, as query sequences. We further conducted tblastn of blast v2.11.0 [77] searches using these query sequences against the whole genomes of closely related species, with search parameters set to *-evalue 1e-5 -num_threads 16*. For the potential loci obtained from the tblastn searches, we extended 2000 bp upstream and downstream of each locus. Subsequently, we performed detailed predictions using GeneWise v3 (RRID:SCR_015054) [78], which allowed us to accurately identify the coding regions of coding genes. Finally, we compared the predicted proteins against the non-redundant protein databases of human, chicken, and green anole to confirm the reliability of our predictions.

### Gene synteny analysis

Based on the annotation methods previously applied to *P. vitticeps*, we rapidly performed batch predictions of coding genes on the genome assemblies of nine closely related species using GeMoMa. The annotation GFF3 files and coding sequences were collected. Syntenic gene pairs between *P. vitticeps* and related species were identified using the Python version of MCScan, JCVI v1.1.16, with the following command: *python3 -m jcvi.compara.catalog ortholog --score=0.95 -- no_strip_names*. Syntenic blocks were filtered using the command: *python3 -m jcvi.compara.synteny mcscan --iter=1*. We combined all species’ syntenic gene pairs based on *P. vitticeps* loci using the command: *python3 -m jcvi.formats.base join --boheader*. For the synteny plot of each target gene, we selected the target gene and its flanking 10 genes from the *P. vitticeps* locus and generated the plot using the command: *python3 -m jcvi.graphics.synteny*.

### Phylogenetic analysis and gene evolution

The protein sequences of each orthologous group of sex-determining genes were aligned using MAFFT v7.471 (RRID:SCR_011811) [79] with the parameters: mafft --anysymbol --maxiterate 1000 --localpair pep.fa. The multiple alignment sequences were then processed to remove spurious sequences or poorly aligned regions using trimAl v1.4.rev15 (RRID:SCR_017334) [80] with the command: trimal -in pep.fa.aln -out pep.fa.aln.trimal -gt 1. Maximum likelihood inference of phylogenetic relationships was performed using IQ-TREE v2.2.0.3 (RRID_SCR_017254) [81] with the parameters: iq-tree -m TEST -bb 1000 -bnni. The ratios of non-synonymous to synonymous substitutions within six sex-determining gene pairs were calculated using KaKs_Calculator v2.0 (RRID:SCR_022068) [82] with its default settings.

### Transcriptome source across different sexes and developmental stages

For the male and female gonadal transcriptome data, we used previously published datasets, which included *P. vitticeps* ZZ and ZW individuals at four developmental stages (6/7, 12, 16, and adult) [14, 46]. The early-stage gonadal RNA-seq source accession numbers are SRP304423. The adult source accession numbers used for testes of ZZ male are ERR753529 and ERR413070, and for ovaries of ZW female are ERR753530 and ERR413082.

### RNA-seq and differential expression analysis

Paired-end raw reads underwent adapter trimming using SOAPnuke v1.5.6 (RRID: SCR_015025) [56] with *filter -n 0.03 −l 20 -q 0.3 -p 1 -Q 2 -G −5 1 -t 10,0,10,0 -E 70*. Quality control of the sequencing reads was conducted using FastQC v0.12.1 (RRID:SCR_014583). The clean data were mapped to genome using STAR v2.7.11a (RRID:SCR_004463) [83]. Counts of reads mapping to genes were obtained using FeatureCounts v2.0.1 (RRID:SCR_012919) [75] against the top-length isoform annotation with the following command: -T 16 -g gene_id -t exon. To investigate gene expression changes from stage 6/7 to adult, differential gene expression analysis was conducted between these stages for each sex, ZZ and ZW. Differential expression was performed in R v4.0.2 using the DESeq2 package v1.28.1 (RRID:SCR_015687) [84]. Additionally, we processed the raw expression matrix using TPM (Transcripts Per Million) counts to ensure comparability between samples, and ultimately used these values to generate histograms of gene expression.

## Additional Files

**Supplementary Fig. S1.** Sex identification and additional description of genome assembly.

**Supplementary Fig. S2.** Quality validation of the genome assembly.

**Supplementary Fig. S3.** Analysis of missing sequences and related genomic features.

**Supplementary Fig. S4.** Expression matrix of 15,758 lncRNAs across 8 different tissues.

**Supplementary Fig. S5.** Analysis of sex chromosome.

**Supplementary Fig. S6.** Analysis of SDR genes in *P. vitticeps* and related species

**Supplementary Table S1.** Summary statistics of sequencing data for the genome assembly and annotation of *Pogona vitticeps*.

**Supplementary Table S2.** Detailed statistics for CycloneSEQ long-read RNA-seq data from eight tissues.

**Supplementary Table S3.** Detailed statistics for DNBSEQ short-read RNA-seq data from eight tissues.

**Supplementary Table S4.** Estimation of genome size and heterozygosity of *Pogona vitticeps* by *k*-mer analysis.

**Supplementary Table S5.** Improvement in continuity and completeness of genome assembly generated by each of the seven assembly steps as stated in main text.

**Supplementary Table S6.** Genome features between Pvit2024 and Pvit2015.

**Supplementary Table S7.** Summary of comprehensive assessment of genome completeness for 67 long-read sequenced Squamata species using BUSCO and Compleasm.

**Supplementary Table S8.** Summary of telomere and centromere location information of *Pogona vitticeps* genome.

**Supplementary Table S9.** Comparison of telomeric region characteristics between *Pogona vitticeps* assembly and 28 squamate species whose genome reached chromosome level.

**Supplementary Table S10.** Annotation of repeat sequences.

**Supplementary Table S11.** Gene model and function annotation.

**Supplementary Table S12.** 47 protein-coding genes and 65 lncRNA genes that are missing or unknown to be Z-linked in prior Pvit2015 assembly.

**Supplementary Table S13.** Data information for identification of S0 region.

**Supplementary Table S14.** Overview of all sequenced embryonic gonad samples included in this study.

**Supplementary Table S15.** Differential expression profiles of Z-link genes across various developmental stages.

**Supplementary Table S16.** Quality control of long-range sequencing data from two platforms.

## Abbreviations

LncRNA: Long non-coding RNA
PAR: pseudo-autosomal region
SDR: sexually differentiated region
EBP: Earth BioGenome Project
VGP: Vertebrate Genomes Project
WGS: whole genome sequencing
PCR: polymerase chain reaction
HMW: high molecular weight
TRUs: telomeric repeat units
T2T: telomere-to-telomere
BUSCO: Benchmarking Universal Single-Copy Orthologs
QV: consensus quality value
RNA-seq: RNA-sequencing
TPM: Transcripts Per Million
NCBI: The National Center for Biotechnology information
UTR: Untranslated Region
5’-UTRs: five prime untranslated regions
3’-UTR: three prime untranslated regions
A3: alternative 3’ splice site
A5: alternative 5’ splice site
AF: alternative first exon
AL: alternative last exon
MX: mutually exclusive exons
RI: retained intron
SE: skipped exon
GSD: genetic sex determination
chr2qter: the terminal region of the long arm of chromosome 2
TSD: Temperature-dependent sex determination
TSO: template switch oligo sequence
RTP: reverse transcription primer
NOR: nucleolus organizer region
tRNAs: transfer RNAs
rRNAs: ribosomal RNAs.

## Code availability

The bioinformatic tools used in this work, including version, setting and parameters, had been described in the Methods section. Default parameters were applied if no parameters were mentioned for a tool. The pipeline for genome *de novo* assembly, annotation, and chromosome contact analysis, and the custom code and scripts for sex chromosomes analysis in this work had been deposited at GitHub (https://github.com/guoqunfei/Pvit_T2T). All figures were plotted in R.

## Data Availability

CycloneSEQ long-read WGS data, DNBSEQ short-read WGS data, CycloneSEQ long-read RNA-seq data, DNBSEQ short-read RNA-seq data, CycloneSEQ based Pore-C data, and Hi-C data in this study are deposited in NCBI Sequence Read Archive (SRA) under BioProject accession no. PRJNAxxxxxx and in the CNGB Nucleotide Sequence Archive (CNSA) of China National Gene Bank DataBase (CNGBdb) under accession no. CNP0005509. Genome assembly, annotation, other supporting data, and material are available in the *GigaScience* GigaDB database.

## Authors’ contributions

Q.L., Y.D., and Y.G. conceived the study. W.D., W.J., and Y.W. performed sample collection and tissue dissection. F.G., T.Z., X.S., and X.Y. conducted the CycloneSEQ long-read WGS experiments and sequencing. J.C., C.C., and J.Y. performed CycloneSEQ based Pore-C experiments and sequencing. Q.G., Y.P., W.C., H.C., Y.Z., and C.Y. performed genome assembly and annotation. Q.G., Y.P., Y.M., Y.Z., S.S., and B.W. conducted comparative analyses, combined the results, and draw figures. X.X. contributed computing resources. Q.L. wrote the manuscript with the inputs from all authors. All authors read and approved the final manuscript.

## Acknowledgements

We thank Professor Arthur Georges and Dr. Sarah L. Whiteley from the University of Canberra for their providing transcriptome data of *P. vitticeps* across early developmental stages. We also thank the China National GeneBank for providing computing resources.

## Supplementary Figures

**Figure S1:**
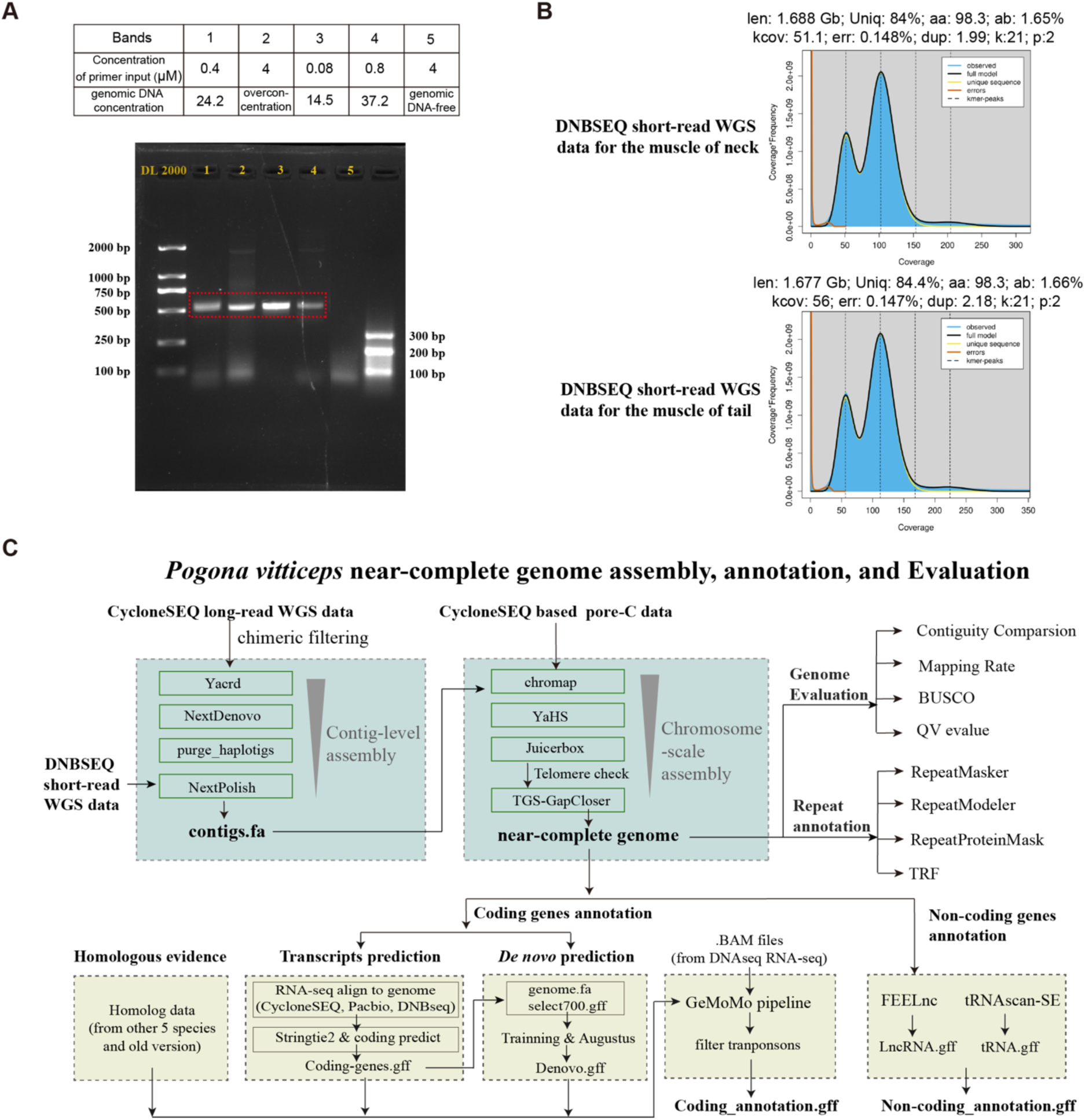
Sex identification and additional description of genome assembly (related to Fig. 1). (A) Amplification of the DNA fragment of the PCR primers that flank two W-chromosome-specific deletions. Band 1 to 4 were made using different concentrations of primers and genomic DNA. The fifth band without genomic DNA was the control group. (B) The 21-mer frequency distribution of two libraries of DNBSEQ short-read WGS data. (C) Comprehensive assembly and annotation workflow for the near-complete genome of *P. vitticeps*.

**Figure S2:**
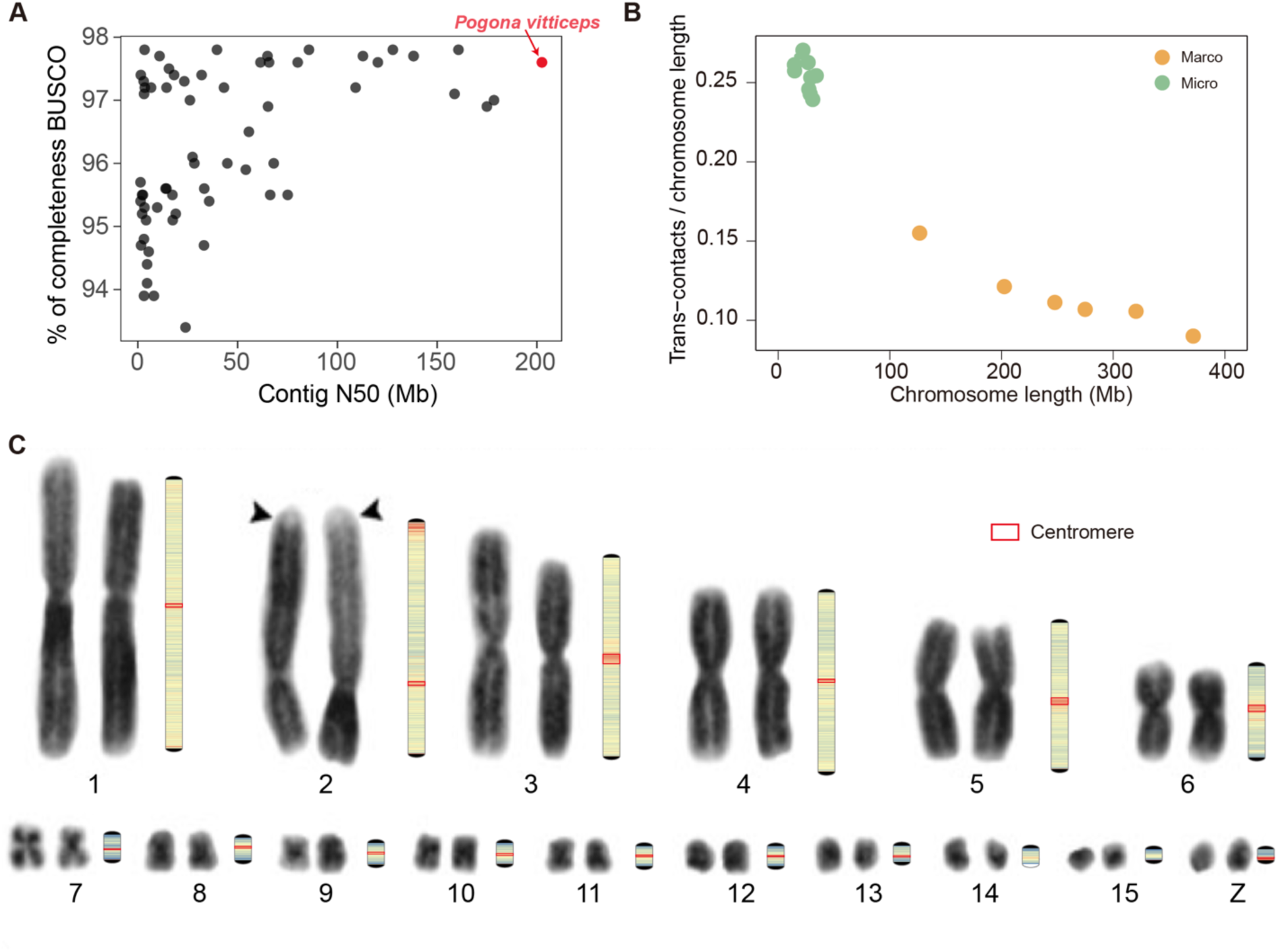
Quality validation of the genome assembly (related to Fig. 2). (A) The comparison of BUSCO complete score based on 7,480 Sauropsida conserved genes and contig N50 between Pvit2024 and 66 publicly available squamate genomes assembled by long-read data. (B) Trans-contact for each chromosome were calculated by ∼153X Hi-C_1 data. (C) A high degree of consistency between the location of the centromere region identified by the near-complete genome with the depressed region in the karyotype map from Young et al., which is generally believed to be the centromere.

**Figure S3:**
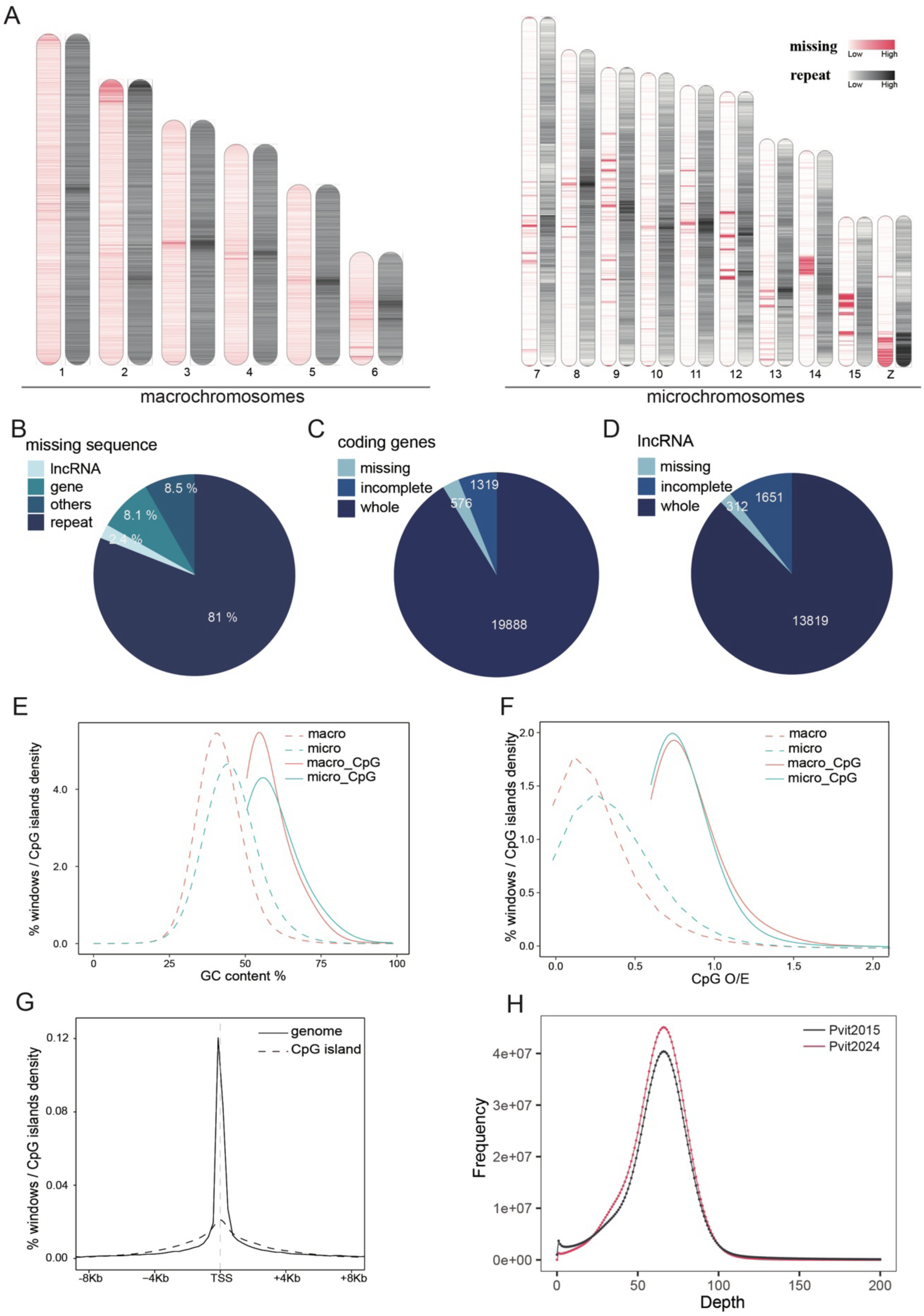
Analysis of missing sequences and related genomic features (related to Fig. 3). (A) Overview of missing and repeat sequences distribution across all chromosomes. The red range indicates missing sequences, and the black range represents repeat sequences. (B) Proportion of various types of elements within the missing sequences. (C) Types and proportions of missing coding genes in the Pvit2015. (D) Types and proportions of missing lncRNAs in the Pvit2015. (E) In the genome, use a non-overlapping sliding window method to scan the genome, and the window size was set to 200bp. By calculating the GC content and CpG O/E value of each window, windows that satisfied the GC content ≥ 50% and CpG O/E ≥ 0.6 were identified as CpG island. The GC content density distribution maps of all windows and CpG islands in the genome were obtained, in which macro- and micro-chromosomes were displayed separately due to different genomic characteristics. (F) CpG O/E value density distribution of all windows and CpG islands in the genome, macrochromosomes and microchromosomes are displayed separately. (G) According to the genome annotation file, TSS (transcription start site) was located, and the CpG island closest to each TSS was found within 8 kb upstream and downstream, and its relative distance was calculated. For comparative analysis, 200 bp windows equal to the CpG islands were randomly selected in the genome, and the closest relative distance between these random windows and each TSS were also counted. The density distribution maps of the closest distances between TSS and the random windows of the genome and CpG islands were obtained to compare the distribution characteristics. (H) Distribution of base coverage depth after mapping DNBSEQ reads to the Pvit2015 and Pvit2024 genomes.

**Figure S4:**
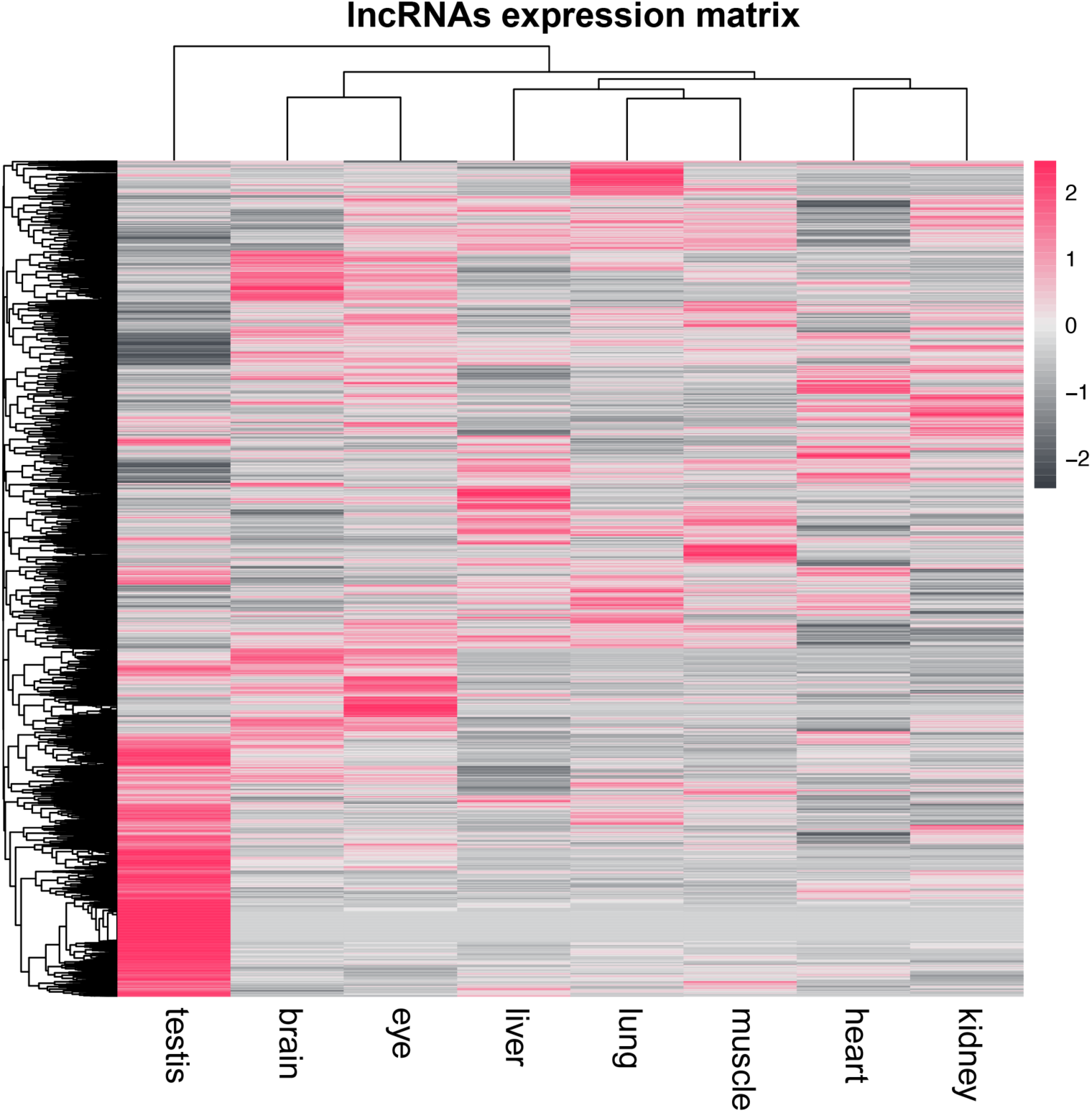
Expression matrix of 15,758 lncRNAs across 8 different tissues (related to Fig. 4).

**Figure S5:**
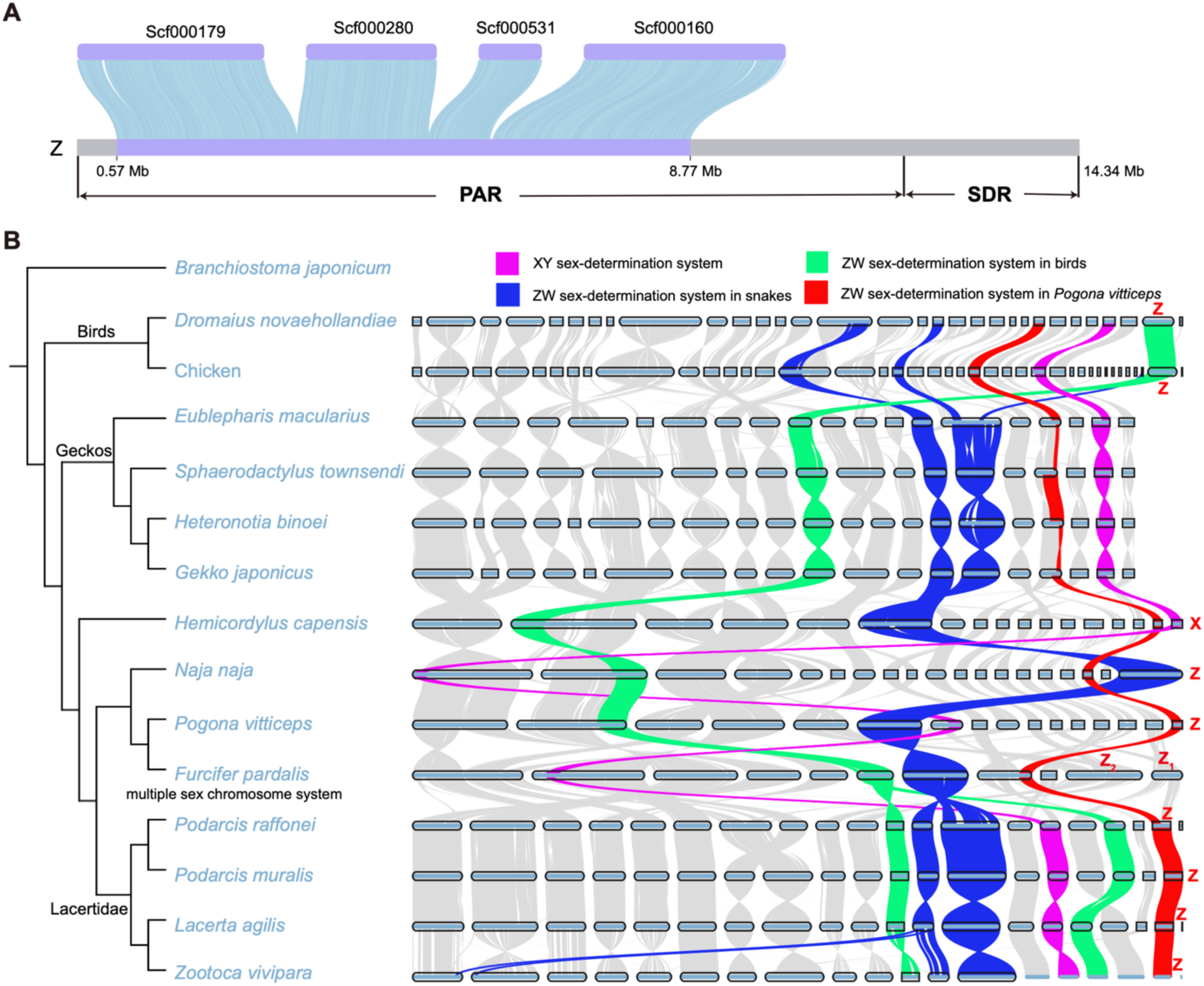
Analysis of sex chromosome (related to Fig. 5). (A) Collinearity between the four known Z-linked scaffolds identified by Deakin et al. and the Z chromosome of Pvit2024. (B) The construction of sex chromosome homologous regions between the identified sex chromosomes of squamate species and the Z chromosomes of two closely related species, the chicken (*Gallus gallus*) and the emu (*Dromaius novaehollandiae*), based on comprehensive gene collinearity analysis.

**Figure S6:**
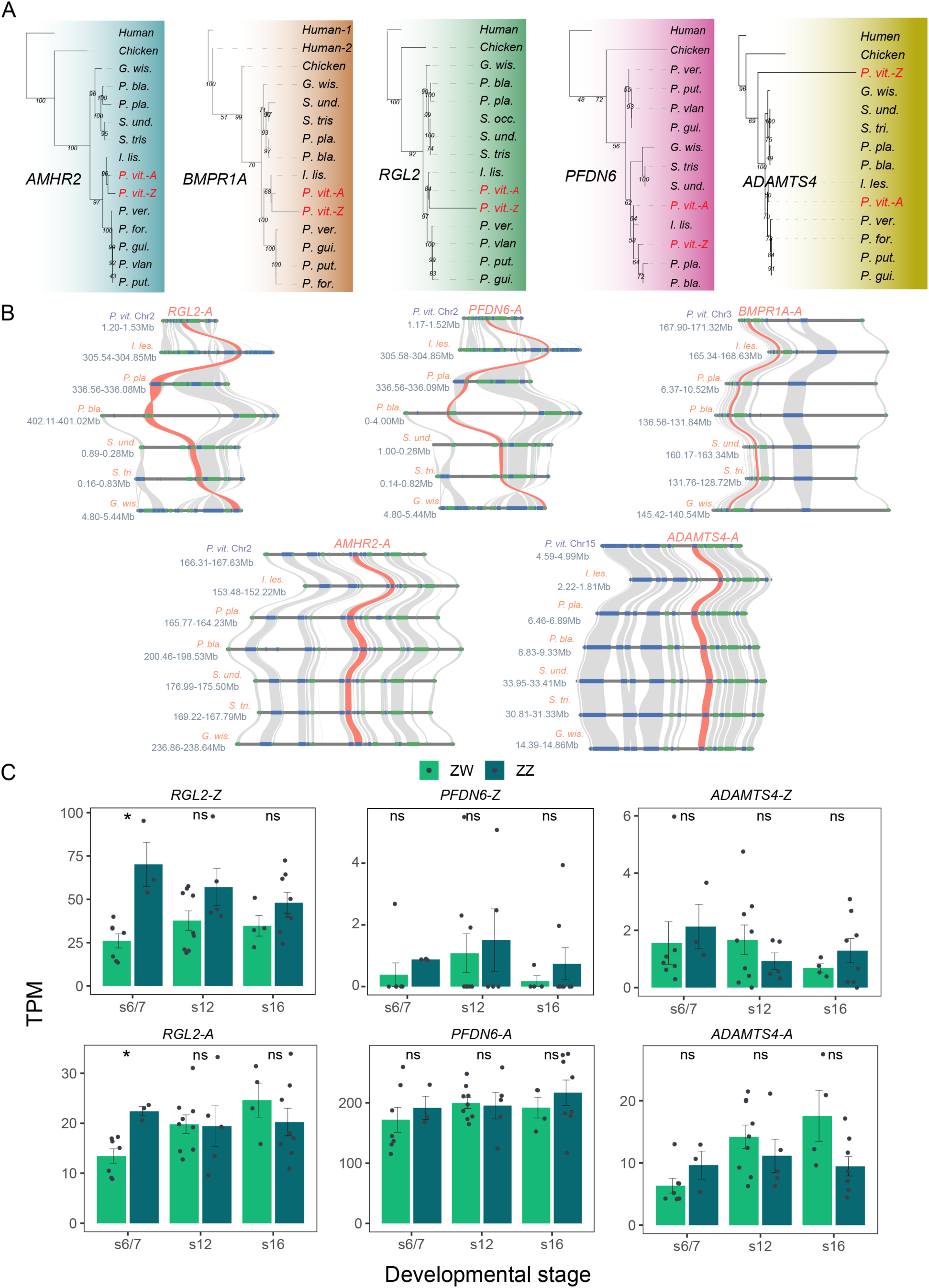
Analysis of SDR genes in *P. vitticeps* (related to Fig. 6). (A) Phylogenetic relationships of five genes in different species. Light red font indicates the two copies of SDR-related genes in *P. vitticeps*. (B) Synteny relationships of autosome SDR genes and their flanking 10 genes in closely related species. The target genes are highlighted in orange, and the surrounding genes are shown in grey. (C) Differential expression of SDR gene pairs between ZZ and ZW during different developmental stages. Benjamini-Hochberg analysis of variance: ns (not significant), \**P*<0.05, \*\**P*<0.01, \*\*\**P*<0.001.

